# Null Models for Community Detection in Spatially-Embedded, Temporal Networks

**DOI:** 10.1101/007435

**Authors:** Marta Sarzynska, Elizabeth A. Leicht, Gerardo Chowell, Mason A. Porter

## Abstract

In the study of networks, it is often insightful to use algorithms to determine mesoscale features such as “community structure”, in which densely connected sets of nodes constitute “communities” that have sparse connections to other communities. The most popular way of detecting communities algorithmically is to optimize the quality function known as modularity. When optimizing modularity, one compares the actual connections in a (static or time-dependent) network to the connections obtained from a random-graph ensemble that acts as a null model. The communities are then the sets of nodes that are connected to each other densely relative to what is expected from the null model. Clearly, the process of community detection depends fundamentally on the choice of null model, so it is important to develop and analyze novel null models that take into account appropriate features of the system under study. In this paper, we investigate the effects of using null models that take incorporate spatial information, and we propose a novel null model based on the radiation model of population spread. We also develop novel synthetic spatial benchmark networks in which the connections between entities are based on distance or flux between nodes, and we compare the performance of both static and time-dependent radiation null models to the standard (“Newman-Girvan”) null model for modularity optimization and a recently-proposed gravity null model. In our comparisons, we use both the above synthetic benchmarks and time-dependent correlation networks that we construct using countrywide dengue fever incidence data for Peru. We also evaluate a recently-proposed correlation null model, which was developed specifically for correlation networks that are constructed from time series, on the epidemic-correlation data. Our findings underscore the need to use appropriate generative models for the development of spatial null models for community detection.

**PACS numbers:** 87.19.Xx,89.20.-a,89.75.Fb,05.45.Tp

## I. Introduction

A network formalism is often very useful for describing complex systems of interacting entities [1, 2]. Scholars in a diverse set of disciplines have studied networks for many decades, and network science has experienced particularly explosive growth during the past 20 years [1]. The most traditional network representation is a static graph, in which nodes represent entities and edges represent pairwise connections between nodes. However, many networks are time-dependent [3, 4] or multiplex (include multiple types of connections between nodes) [5, 6]. Moreover, network structure is influenced profoundly by spatial effects [7]. To avoid discarding potentially important information, which can lead to very misleading results, it is thus crucial to develop methods that incorporate features such as time-dependence, multiplexity, and spatial embeddedness in a context-dependent manner [3, 5, 7]. Because of the newfound wealth of available rich data, it has now become possible to validate increasingly complicated network structures and methods using empirical data.

In the present paper, we study a mesoscale network structure known as *community structure*. A “community” is a set of nodes with dense connections among themselves, and with only sparse connections to other communities in a network [8, 9]. Communities arise in numerous applications. For example, social networks typically include dense sets of nodes with common interests or other characteristics [10], networks of legislators often contain dense sets of individuals who vote in similar ways [11], and protein-protein interaction networks include dense sets of nodes that constitute functional units [12]. The algorithmic detection of communities and the subsequent investigation of both their aggregate properties and the properties of their component members can provide novel insights into the relationship between network structure and function (e.g., functional groupings of newly discovered proteins [13]).

Myriad community detection methods have been developed [8, 9]. The most popular family of methods entails the optimization of a quality function known as *modularity* [14, 15]. To optimize modularity, one compares the actual network structure to some *null model*, which quantifies what it means for a pair of nodes to be connected “at random”. Traditionally, most studies have randomized only network structure (while preserving some structural properties) and not incorporated other features (such as spatial or other information). The standard null model for modularity optimization is the “Newman-Girvan” (NG) null model, in which one randomizes edge weights such that the expected strength distribution is preserved [14, 15]. It is thus related to the classical configuration model [1]. It has become very popular due to its simplicity and effectiveness, and it has been derived systematically through the consideration of Laplacian dynamics on networks [16]. However, it is also a naive choice, as it does not incorporate domain-specific information. The choice of a null model is an important consideration because (1) it can have a significant effect on the community structure obtained via optimization of a quality function, and (2) it changes the interpretation of communities [17–19]. The best choice for a null model depends on both one’s data set and scientific question. In the present paper, we explore the issue of null model choice in detail in the context of spatially embedded and temporal networks.

Most existing research on community detection does not incorporate metadata about nodes (or edges) or information about the timing and location of interactions between nodes. However, with the increasing wealth of space-resolved and time-resolved data sets, it is important to develop community detection techniques that take advantage of the additional spatial and temporal information (and of domain-specific information, such as generative models for human interactions [20]). Indeed, community detection in temporal networks has become increasingly popular [21–27], but the majority of methods use networks that are constructed from either static snapshots of data or aggregations of data over time windows. Few investigations of community structure in temporal networks have used methods that exploit temporal structure (see, e.g., [24, 27]). There is also starting to be more work on the influence of space on community structure [20, 28–31], but much more research is necessary.

In the present paper, we use modularity maximization to study communities in spatially embedded and time-dependent networks. We compare the results of community detection using two different spatial null models — a *gravity null model* [20] and a new *radiation null model* — to the standard NG null model using novel synthetic benchmark networks that incorporate spatial effects via distance decay or disease flux as well as temporal correlation networks that we constructed using time-series data of recurrent epidemic outbreaks in Peru. We also evaluate a recently-proposed *correlation null model*, which was developed specifically for correlation networks that are constructed from time series [18], on the epidemic-correlation data.

Network methods have become increasingly prevalent in the modeling of infectious diseases [32]. Most studies focus on the importance of interpersonal contact networks on the disease spread on an individual level. Our direct analysis of disease data in the present paper provides a complementary (e.g., more systemic) approach. Our work also complements other approaches, such as large-scale compartmental models that incorporate transportation networks to link local populations. Such models have been used to study large-scale spatial disease spread (e.g., to examine the influence of features such as spatial location, climate, and facility of transportation on phenomena such as disease persistence and synchronization of disease spread) [7, 33–35].

The rest of the present paper is organized as follows. In Section II, we give an overview of networks and community detection. We also discuss the gravity null model and introduce a new radiation null model. We give our results for synthetic spatial networks in Section III, and we give our results for correlation networks that we construct from disease data in Section IV. We summarize our results in Section V. In appendices, we include the results of additional numerical experiments from varying parameters in the benchmark networks. We also include an additional examination of the similarity between network partitions for the benchmarks and the dengue fever correlation networks.

## II. Networks and Community Structure

A network describes a set of entities (called *nodes*) that are connected by pairwise relationships (called *edges*). In the present paper, we study weighted networks which are *spatially embedded*: each node represents a location in space. One can represent a weighted network with *N* nodes as an *N × N* adjacency matrix *W*, where an edge *W_ij_* represents the strength of the relationship between nodes *i* and *j*. We seek to find *communities*, which are sets of nodes that are densely connected to each other but sparsely connected to other dense sets in a network [8, 9].

We wish to study the evolution of network structure through time. The simplest way to represent temporal data is through an ordered set of *static networks*, which can arise either as snapshots at different points in time or as a sequence of aggregations over consecutive time windows (which one can take either as overlapping or nonoverlapping). Static networks provide a good starting point for the development and investigation of new methods — which, in our case, entails how to incorporate spatial information into null models for community detection via modularity maximization. However, they do not take full advantage of temporal information in data that changes in time. For example, it can be hard to track the identity of communities in temporal sequences of networks [24].

To mitigate the community-tracking problem, we also use a type of *multilayer network* [5, 6] known as a multislice network [24]. This gives an *N × N × m* adjacency tensor 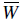 that has *m* layers and *N* nodes in each layer. The intralayer edges in the network are exactly the same as they were for the sequence of static networks. The tensor element 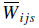 gives the weight of an intralayer edge between nodes *i* and *j* in layer *s*. Additionally, each layer has a copy of node *i*, and it is connected to itself in consecutive layers *s* and *r* using interlayer edges of weight *C_isr_*. In this paper, we will suppose for simplicity that *C_isr_ = ω* ∈ [0, ∞), but one can also consider more general situations [5, 36]. A multislice network can have up to (*N* × *m*) *multilayer nodes* (i.e., node-layer tuples), each of which corresponds to a specific (node, time) pair. Hence, this structure makes it possible to detect temporally evolving communities in a natural way.

For our computations of community structure, we flatten the *N* × *N* × *m* adjacency tensor into a (*N* × *m*) × (*N* × *m*) adjacency matrix, such that the intralayer connections are on the main block diagonal and the interlayer connections occur on the off-block-diagonal entries. We detect communities by maximizing modularity, which we use to describe the “quality” of a particular network partition into communities in terms of its departure from a null model [14]. The null model amounts to a prior belief regarding influences on network structure, so it is important to carefully consider the choice of null model [18, 20, 27].

For a weighted static network *W*, modularity is [37]

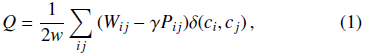

where 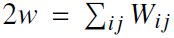 is the total edge weight, *c_i_* denotes the community that contains node *i,* the function *δ* is the Kronecker delta, and *P_ij_* is the *ij*-th element of the null model matrix. One can examine different scales of community structure by incorporating a resolution parameter *γ* [38, 39]. Smaller values of *γ* tend to yield larger communities and vice versa.

For multislice networks, modularity is given by

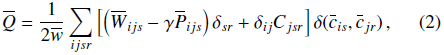

where 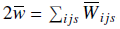, the quantity 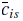 denotes the community that contains node *i* in layer *s,* and 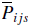 is the *ij*-th element of the null model tensor in layer *s* [24].

To detect communities via modularity maximization, one searches the possible network partitions for the one with the highest modularity score. Because exhaustive search over all possible partitions is computationally intractable [40], practical algorithms invariably use approximate optimization methods (e.g., greedy algorithms, simulated annealing, or spectral optimization), and different approaches offer different balances between speed and accuracy [8, 9].

In the present paper, we optimize modularity using a two-phase iterative procedure similar to the Louvain method [41]. However, rather than using the adjacency matrix *W*, we work with the modularity matrix *B* with elements 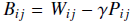 for static networks and with the modularity tensor with elements 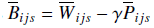 for multislice networks [42].

The employed Louvain-like algorithm [42] is stochastic, and a modularity landscape for empirical networks typically includes a very large number of nearly-optimal partitions [17]. For each of our numerical experiments, we thus apply the computational heuristic 50 times to obtain a *consensus community structure* [43] by constructing an *association matrix A*^rep^ (where the entries 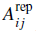 represent the fraction of times that nodes *i* and *j* are classified together in the 50 partitions) and performing community detection on *A*^rep^ using the uniform null model 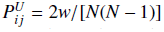 [27]. We choose the uniform null model in order to detect the strongest community structure in the association matrix (i.e., one that is often detected by the original optimization process).

Because community detection using modularity maximization is strongly parameter-dependent, one might also sometimes be interested in the most *persistent communities* across a range of values of the resolution parameter *γ* or (for multislice networks) the interlayer edge weight *ω* [12, 27]. To obtain these, we construct an association matrix *A*^persist^ across a range of parameter values (where the entries *A_ij_^present^* represent the fraction of times that nodes *i* and *j* are classified together in network partitions across different parameter values) and perform community detection on *A*^persist^ using the uniform null model to detect the most persistent community structure in the association matrix (i.e., one that is often detected by the original optimization process).

For multislice networks, we perform community detection and then consensus clustering using the same basic procedure. This yields an assignment of each multilayer node (i.e., node-layer tuple) to a community. We are also sometimes interested in community assignments of the original entities (i.e., a partition of the set of nodes regardless of what layer they are in). For example, we might wish to compare the result of algorithmic community detection to known partitions, such grouping a node (i.e., province) by climate, population, administrative region, etc. To do this, we perform what we call *province-level community detection*, which proceeds in two rounds: (1) we detect communities in a multislice network using any method and null model of choice; (2) we use this partition to construct an *N × N* province-level association matrix (i.e., a matrix *A*^province^ where entries *A_ij_*^province^ represent the fraction of times that nodes *i* and *j* are classified together in all layers), and we detect province-level communities by maximizing modularity on this association matrix using a uniform null model. We choose the uniform null model to detect the most temporally persistent community structure in the association matrix (i.e., one that is often detected in multiple layers).

### A. Null Models for Community Detection

The choice of null model is vital for the detection of communities using modularity maximization [17, 18, 27]. The most common choice is the Newman-Girvan (NG) null model, which randomizes a network such that the expected strength sequence of nodes is preserved [44, 45]. For static, weighted networks, the NG null model is given by

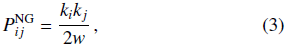

where 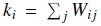 is the strength (i.e., weighted degree) of node *i* and 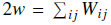 is the total edge weight in the network.

For multislice networks, the NG null model is [24]

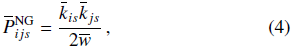

where 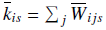 is the intralayer strength of node i in layer *s* and 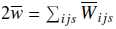.

Despite its popularity and demonstrated effectiveness in many situations, the NG null model is naive in the sense that it does not incorporate problem-specific information (such as spatial embeddedness). It only takes node strengths into account, and consequently it is not suitable for all applications. It is often important to incorporate additional (domain-specific or even problem-specific) information, and what one considers to be connected “at random” depends fundamentally on the research question of interest.

#### 1. Spatial Null Models: Gravity Model

In many spatially embedded networks, proximity has a strong effect on the connections between nodes, as (all else held equal) neighboring nodes are more likely to be connected to each other (and their connections are likely to have to have larger weights) than nodes that are far away [7, 20]. Moreover, proximity can mask other underlying influences. Consequently, incorporating the expected influence of proximity on edge weights into null models for community detection should make it possible to discover new and important types of structures.

Expert et al. [20] proposed a spatial null model that was inspired by the “gravity model” of human mobility [46–49]. A gravity model assumes that the interaction between two locations is proportional to their importance (e.g., population), but it decays with distance.

In the standard gravity model, the interaction between locations *i* and *j* with respective populations *n_i_* and *n_j_* that are a distance *d_ij_* apart is

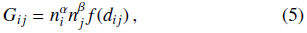

where the “deterrence function” *f*(*d*) describes the effect of space on node interactions. Common choices for the deterrence function include inverse proportionality to distance (i.e., *f*(*d_i j_*) = 1/*d_ij_*), inverse proportionality to squared distance (i.e., 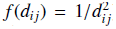), exponential decay (i.e., 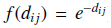), and other interactions of the form *f*(*d_ij_*) *= d_ij_^k^*) [7]. It is common the estimate the parameters *α, β,* and *κ* using regression. Gravity models have been employed successfully during the past half century to model spatial interactions such as population migration [7, 50, 51], trade [52], and disease spread [35].

The simplest form of a gravity-like interaction in Eq. (5), with *α = β =* 1 and *κ =* -1, was incorporated into a *gravity null model* [20], to give

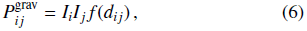

where *I_i_* is the importance of node *i*. One estimates the “deterrence function”

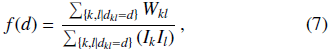

from data for all nodes at distance *d* between them in a data set. Expert et al. [20] used *I_i_* = *n_i_* (where *n_i_* denotes the population of province *i*) as their measure of node importance. After briefly experimenting with variations, such as using population density or a logarithm of the population (i.e., *I_i_* = log(*n_i_*)) and observing no significant differences in performance, we will follow their lead. Another simple choice is node strength (i.e., *I_i_* = *k_i_* = Σ*_j_ W_ij_*), though the null model then becomes very similar to the usual NG null model [20]. Moreover, if *f*(*d*) does not depend on distance, then the null model becomes exactly the NG null model in that case.

In most data sets, distances are continuous, so one needs to bin distance data to obtain enough nodes in each distance bin to construct a meaningful deterrence function *f*(*d*) in Eq. (7). In our calculations, we bin the distances into equal-distance bins (e.g., every *b* km). After examining the effects of bin size on algorithmic community structure — in particular, we studied the effect of bin size on the deterrence function and the effect of bin size on the partition quality and similarity between algorithmic partitions — we select a bin size large enough so that the deterrence function is relatively smooth but small enough that it shows a downward trend. We will give the specific bin sizes for spatial benchmark and dengue correlation networks in their respective Sections. For the benchmark networks we can test the similarity of algorithmic partitions to the planted community structure at different bin sizes.

Alternative binning methods include binning into equal-sized bins (e.g., each bin containing *c* elements). After testing the choice of binning procedure on the benchmark networks before applying the null model to empirical data and observing no qualitative differences in null model performance, we selected the equal-distance method for the rest of the paper.

Combining Eqs. (6) and (7) allows us to write the gravity null model as

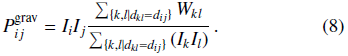

Expert et al. used the null model (8) to uncover a linguistic partition of a network of Belgian mobile phone calls into the French and Flemish speaking parts of Belgium. This partition was obscured by geographical communities when using the NG null model [20].

In the present paper, we generalize the gravity null model to a multislice setting by calculating a separate gravity null model for each layer *s*. The resulting multislice gravity null model is

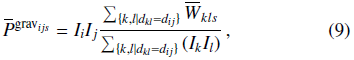

where we have assumed that the population stays constant across time. If one has reliable information about changes in population with time, one can incorporate such information into the null model (9) by substituting *I_i_* with an analogous quantity *I_is_* that depends both on the node *i* and on the layer *s*.

#### 2. Spatial Null Models: Radiation Models

Gravity models include multiple parameters that one needs to either choose arbitrarily or estimate from data. Moreover, by their design, gravity models are unable to predict different fluxes between locations that are the same distance apart but which have regions with different population densities between them. For example, one would expect a higher flux of infectious disease between two locations that are separated by a space with high population density than between locations that are separated by a space with low population density (because of the higher availability of susceptible hosts in the latter case) [53]. By contrast, one would expect a smaller commuting flux between such locations in the latter case due to higher availability of nearby jobs, as this reduces peoples’ willingness to commute for longer distances [54].

A recent model that was developed to attempt to address these issues is the radiation model [54], which was designed for population flows and has subsequently been applied successfully in several situations [55, 56]. Because the radiation model is designed to capture human mobility between populations, and the long-distance spread of many infectious diseases — including dengue — is believed to be largely due to long-distance mobility [57], the radiation model might provide a useful but simplified description for the spread of disease across space. In this section, we use it to construct a new spatial null model for community detection that we believe might be well-suited for studying the long-distance spread of dengue.

The mean commuting flux predicted by the radiation model for locations *i* and *j* with populations *n_i_* and *n_j_* is

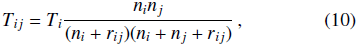

where *r_ij_* is the population between locations *i* and *j*, and *T_i_* is the number of commuters in location *i*. A simple way to calculate *r_ij_* is to use the population *q_ij_* in the circle of radius *d_ij_* centered at *i* and subtract the total of the populations at the origin and destination. That is, *r_ij_* = *q_ij_* − (*n_i_* + *n_j_*). Although the radiation model is relatively recent [54], several modifications to it have already been proposed. These include incorporating a normalization for finite systems [56] and the development of a general framework that includes ideas from the radiation, gravity, and intervening-opportunities models [58].

We propose a novel null model for community detection based on the original formulation of the radiation model [54]. We use a similar formulation to Eq. (8) to incorporate both the expected distance-dependent flux and the actual network structure. To avoid creating a directed network, we use a symmetrized predicted flux

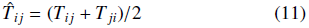

between nodes *i* and *j*. ^1^ We thereby construct the *radiation null model*

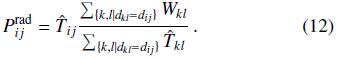

In Section IV, we will study community structure in empirical data from several years of dengue fever occurrences in Peru. Because we do not possess detailed data on the commuting patterns in Peru (see the description of our data in Section IV A), we assume that commuters are distributed uniformly across space. We can then simplify Eq. (10) by substituting *T_i_* = *T_f_n_i_*, where *T_f_* is the fraction of the population that commutes. Because the quantity *T_f_* is present in both the numerator and denominator of Eq. (12), we can now cancel it out. However, if one possesses commuting data, it would be desirable to use it to improve the radiation null model.

We also extend the radiation null model to a multislice setting in an analogous manner to the gravity null model. The multislice radiation null model is

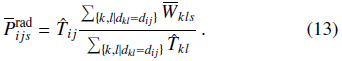

Again, one can incorporate temporal data about population sizes and thereby replace *T_ij_* with *T_ijs_* to improve the null model.

#### 3. Spatial Null Models: Other Models

The incorporation of spatial information into null models for community detection is an important problem, and several other ideas have been proposed recently. For example, Cerina et al. [28] focused on disentangling the correlation between node attributes and space, so they used a simple exponential decay: 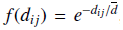, where *d̄* is the mean distance between nodes in a network. Shakarian et al. [30] focused on finding geographically-disperse communities, so they introduced a decay constant *θ* such that 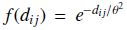. Another recently-proposed null model was used to attempt to find geographically-proximate communities [29].

As the exact nature of the influence of spatial distance on interactions in the dengue fever data is unclear, we decided to focus only on null models that include a contribution from the data, rather than using null models with an arbitrarily chosen functional dependence. Thus, we do not test these null models in the present paper.

## III. Synthetic Benchmark Networks

To test the performance of the spatial null models, we develop novel synthetic benchmark networks that represent idealized spatially-embedded networks with planted community structure.

In what we call the *distance benchmark*, the probability of an edge between two nodes depends only on the geographical distance between nodes and on their community assignments. We assign *N* nodes uniformly at random to positions on the lattice {1, 2,…, *l*} × {1, 2,…,*l*}. We assign a population *n_i_* to each node *i* (which is an idealized “city”). We create two versions of the distance benchmark: the “uniform population distance benchmark” and the “random population distance benchmark”. The uniform population version corresponds to the benchmark in Expert et al. [20]; we assign the same population (*n_i_* = 100) to each node. In the random population benchmark, we assign an integer population uniformly at random from the set {1,…,100}.

We also assign the nodes uniformly at random to one of two communities. In the distance benchmarks, the probability 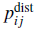 that an edge exists between nodes *i* and *j* at distance *d_ij_* is inversely proportional to distance:

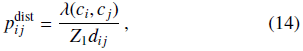

where *c_i_* is the community that contains node *i* and the function *λ*(*c_i_, c_j_*) *=* 1 if nodes i and *j* are in the same community and *λ*(*c_i_, c_j_*) *= λ_d_* otherwise. The “inter-community connectivity” *λ_d_* controls the degree of mixing between communities. When *λ_d_ =* 0, only nodes in the same community are adjacent to each other; when *λ_d_ =* 1, there are no distinct communities. The normalization constant *Z*_1_ ensures that 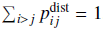. We place *L = μN(N -* 1)/2 edges, where there is an edge between nodes *i* and *j* with probability 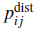 each, and the parameter *μ* ≥ 0 determines the network’s edge density. We interpret multiple edges as weights. We normalize the weights in the network to [0, 1] by dividing each entry by the maximum weight. When we generalize the above benchmarks to a multilayer setting, we thereby yield synthetic multilayer benchmark networks in which the relative magnitudes of interlayer edges and intralayer edges are comparable to those in the disease-correlation networks.

With our *flux benchmark*, we aim to mimic the spread of disease on a network. We allocate its edge weights depending on the mean flux between pairs of nodes that is predicted by the radiation model. We place *N* nodes uniformly at random on the lattice {1, 2, …, *l*} × {1, 2,…,*l*}, and we assign populations and communities in the same manner as for the distance benchmark. Again as with the distance benchmark, we consider both uniform-population and random-population versions of the flux benchmark. Now, however, the edge probability 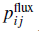 is directly proportional to the predicted radiation-model flux between nodes *i* and *j* (which is turn is inversely proportional to distance *d_ij_*):

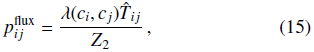

where 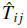 is the mean flux between nodes *i* and *j* that is predicted by the radiation model (see Eq. 10 and 11), and Z_2_ is a normalization constant to ensure that 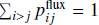.

In Table I, we summarize the four synthetic benchmark networks that we have just introduced.

**TABLE I.**
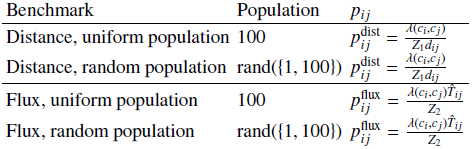
Primary characteristics (i.e., population and edge probability) for the distance and flux benchmarks for static networks. The quantity rand({*a, b*}) signifies that select a number uniformly at random from the set {*a, a* + 1,…, *b*}. Additionally, *λ*(*c_i_*, *c_j_*) *=* 1 if nodes *c_i_* and *c_j_* are in the same community and *λ*(*c_i_, c_j_*) = *λ_d_* otherwise, *d_ij_* is the distance between nodes *i* and *j* in space, and *Z*_1_ and *Z*_2_ are normalization constants.

| Benchmark | Population | $p_{ij}$ |
| --- | --- | --- |
| Distance, uniform population | 100 | $p_{ij}^{\text{dist}} = \frac{\lambda(c_i, c_j)}{Z_1 d_{ij}}$ |
| Distance, random population | $\text{rand}(\{1, 100\})$ | $p_{ij}^{\text{dist}} = \frac{\lambda(c_i, c_j)}{Z_1 d_{ij}}$ |
| Flux, uniform population | 100 | $p_{ij}^{\text{flux}} = \frac{\lambda(c_i, c_j) \hat{T}_{ij}}{Z_2}$ |
| Flux, random population | $\text{rand}(\{1, 100\})$ | $p_{ij}^{\text{flux}} = \frac{\lambda(c_i, c_j) \hat{T}_{ij}}{Z_2}$ |

We create both static (i.e., single-layer) and multilayer benchmarks networks. The static benchmarks enable us to study the performance of modularity maximization using a chosen null model in a simple setting without the additional complications of a multilayer network. However, the multilayer benchmarks are ultimately more appropriate for disease data because they can incorporate temporal evolution.

We begin by placing nodes in space and assigning populations in the same manner as for the static benchmarks. We then assign nodes uniformly at random into one of two communities, and we extend this structure into a multilayer planted community structure with *m* layers. For the “temporally stable” benchmarks, the planted community structure is the same for each layer. For the “temporally evolving” multilayer benchmarks, we change the community assignment of a fraction *p* of the nodes. For each of these nodes, we select a new community assignment uniformly at random, and we change the community of the node in each layer; we start at a layer that we select uniformly at random, and we also change the assignment (to the same new community) in all remaining layers.

We then generate the edges for each layer independently, in the same manner as we generate a static benchmark and using identical parameter values (*N, l, μ, λd*) for each; see Fig. 1. Independent generation of each layer based on the same starting conditions represents differences between observations due to noise and experimental variation.

**FIG. 1.**
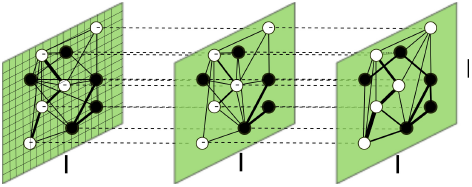
Construction of temporally stable multilayer spatial benchmarks. We assign *N* nodes uniformly at random to positions on a *l×l* lattice (which we show in layer 1) and divide them into two equal-sized communities (black and white) whose nodes we choose uniformly at random. Node *i* has a population of *n_i_*, and each slice has the same set of nodes. For each slice, we allocate edges uniformly at random according to a probability distribution that depends on the type of benchmark; for details, see the text and Table I. We interpret multiple edges as weights, and we visualize these weights using edge thickness. We connect copies of nodes in adjacent slices with interlayer edges of weight *ω* (dashed lines).

For each of the above types of multilayer benchmarks, we set the value of the interlayer edges between corresponding nodes in consecutive layers to be *ω*. Each of the reported community detection results for these benchmarks is an average over consensus community detection (over 50 repeats) for 50 independently drawn instances of a benchmark with the same values of the same parameter values (*N, l, μ, λ_d_*), (*γ, ω*), and (when relevant) *p*.

We evaluate the performance of the NG, gravity, and radiation null models on our benchmarks by comparing algorithmic partitions with the planted community structure using normalized mutual information (NMI) [60]. NMI is an information-theoretic similarity measure that is relatively sensitive to small differences in partitions, such as the move of a single node from one community to another, compared to pair-counting measures such as the Rand coefficient and *z*-Rand scores [61]. This sensitivity makes it suitable for assessing performance on benchmarks that are based on well-defined, ground-truth planted partitions.

NMI is one of many normalized versions of mutual information (MI) [62]. Both MI and NMI are based on the concept of *information entropy*, which is a measure of uncertainty. MI measures the amount of information that one can predict about one random variable (which in the present paper is a partition of a network into communities) based on another one. For a partition *X* = {*X*_1_*, X*_2_*,… X_K_*} with *K* communities and a partition *Y* = {*Y*_1_*, Y*_2_*, … Y_L_*} with *L* communities, MI is defined as

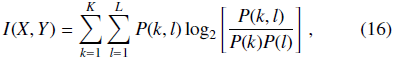

where *P*(*k*) and *P*(*l*) are the marginal probabilities of observing communities *k* and *l* in partitions *X* and *Y*, respectively, and *P*(*k, l*) is the joint probability of observing communities *k* and *l* simultaneously in partitions *X* and *Y*. MI takes values between 0 and min{*H*(*X*), *H*(*Y*)}, where 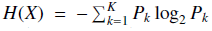 is the entropy of *X*.

Normalized mutual information (NMI) [60] is defined as

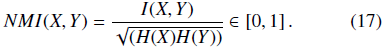

The normalization to lie within the range [0, 1] facilitates interpretation and comparisons. We make use of NMI in the following sections, and we obtain the same qualitative conclusions using variation of information [63], which is a different measure of similarity. See Appendix A for our comparisons using VI.

### A. Results on Static Benchmarks

To emphasize the difference between the gravity and radiation null models, we take *N* = 50 and *l* = 10 to obtain a relatively densely filled lattice. (See Appendix B for the results for a synthetic network with parameter values *N* = 10 and *N* = 90.) We first compare this benchmark versus a situation with parameter values *N* = 100 and *l* = 100 (which are the parameter values that were used in Expert et al. [20]). We test varying bin sizes in uniformly-spaced bins using the parameter values *b* ∈ {10^-4^, 10^-3^, 10^-2^, 0.1} U {1, 2,…, 10}, *l* = 10 and *b* ∈ {1, 2, …, 100}, *l* = 100. We find that bin width makes a large difference on both benchmarks: *b* = 1 produces the highest NMI scores (i.e., it has the “best performance”) and increasing bin width leads to a decrease in performance of both spatial null models (see Fig. 2). This effect is especially pronounced for the gravity null model.

**FIG. 2.**
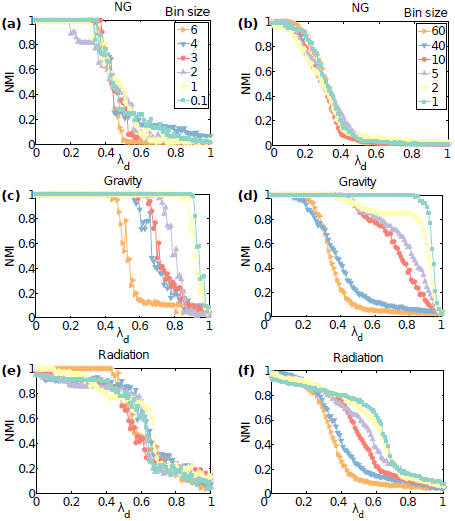
Uniform population static benchmarks: Normalized mutual information (NMI) scores between algorithmically detected and planted community structures in static uniform population distance benchmarks for (left) *l* = 10, *N* = 50 and (right) *l* = 100, *N* = 100, edge density parameter *μ =* 100 and uniform populations of 100 for different bin sizes (colored curves). We detect communities by optimizing modularity using the (top) NG, (middle) gravity, and (bottom) radiation null models.

In both cases, the best aggregate performance of the spatial null models at optimal bin sizes is similar for *l* = 10 and *l* = 100, so we henceforth use the *l* = 10 benchmark with *b* = 1 to lower computational time and memory usage. However, one needs to keep the strong influence of bin size on algorithm results in mind for applications.

We then study the performance of the three null models using several values of the resolution parameter *γ* ∈ {0.5, 0.75, 1, 1.25, 1.5} and the inter-community connectivity *λ_d_* ∈ {0, 0.01,…, 0.99, 1} on static benchmarks with *N =* 50 nodes and lattice size parameter *l* = 10. Smaller values of *γ* tend to yield larger communities and vice versa. Considering larger *λ_d_* increases the level of mixing between the communities and makes community detection more difficult. For simplicity, we fix the density parameter *μ* = 100. As we discuss in Appendix C, the value of *μ* has little effect on the results of community detection when it is above a certain minimum.

For the uniform population distance benchmark, the only factor that influences edge placement is the distance between nodes. On this benchmark, the gravity null model has the best performance, as it is able to find the correct partitions for *λ_d_* 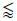 0.82 (see Fig. 3). The radiation null model has the second best performance and is able to find partially meaningful partitions for *λ_d_* 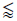 0.74, above which we observe a plateau of “near-singleton” partitions in which most nodes are placed into singleton communities. (We use the term “singleton partition” to refer to a partition in which every node is assigned to its own community.) The NG null model, which does not incorporate spatial information, does much worse than either of the spatial null models; it suffers a sharp decline in performance at *λ_d_* ≈ 0.4. This demonstrates that, although incorporating spatial influence is beneficial for its own sake, we see that using a null model that incorporates population information to study community structure in networks whose structure does not depend on population decreases the performance of community detection. That is, incorporating spatial information is important, but it needs to be done intelligently.

**FIG. 3.**
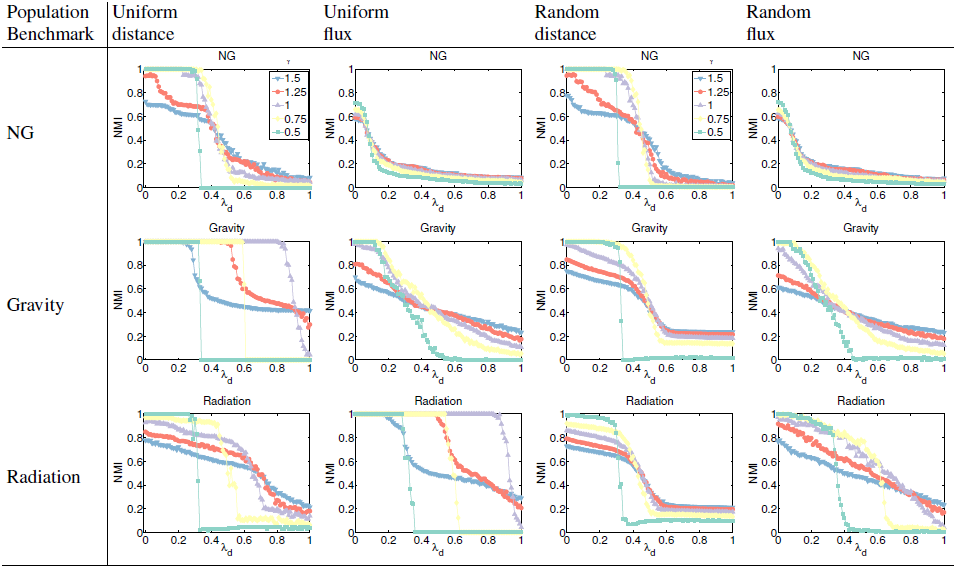
Static benchmarks: NMI scores between algorithmically detected and planted community structures in static benchmarks with *l* = 10, *N =* 50, *μ =* 100 and (columns 1, 2) uniform populations of *n_i_ =* 100 or (columns 3, 4) populations *n_i_* determined uniformly at random from the set {0,…, 100}. We plot NMI for different values of the resolution parameter *γ* (colored curves) as a function of inter-community connectivity *λ_d_* ∈ [0, 1]. We examine both distance benchmarks (in columns 1, 3) and flux benchmarks (in columns 2, 4). We detect communities by optimizing modularity using the (top) NG, (middle) gravity, and (bottom) radiation null models.

On the uniform population flux benchmark — in which we include the population density in the region between two nodes in the flux prediction (so the population density influences edge structure) — the radiation null model outperforms the other null models. The gravity null model comes in second place, and the NG null model is a distant third.

For the random population distance benchmark, we observe a fast deterioration in quality of the detected communities for *λ_d_* 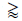 0.4 for all null models, and all null models reach a “near-singleton” regime by *λ_d_* ≈ 0.6. The NG null model has the best performance among the three null models for *λ_d_* 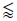. For *λ_d_* 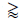 0.43, the gravity null model has the best performance, although the partitions consist largely of singletons for *λ_d_* 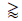 0.6.

For the random population flux benchmark, the radiation null model has the best performance of the three null models. It has the slowest decrease in NMI scores with the increase in *λ_d_*. The gravity null model has the second-best performance, and NG fails even when there is no mixing between the two communities (see Fig. 3). However, even the best performance is much worse on random population benchmarks than it is on the uniform population benchmarks. Note additionally that including population information into the edge placement probability by taking 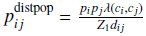 (“distance and population benchmark”) brings back the advantage for the gravity null model (see Appendix D).

Among the parameter values that we consider (*γ* ∈ {0.5, 0.75, 1, 1.25, 1.5}), *γ* = 1 appears to give the best results (i.e., the largest NMI scores). In the near-singleton regime, *γ* = 1.5 outperforms it slightly (see Fig. 3), however this partition is vastly different from the planted partition.

### B. Results on Multilayer Benchmarks

We now study the influence of the resolution parameters *γ* and *ω* on the community quality of multilayer benchmarks networks. We first compare our results to our findings from static benchmarks by varying *γ* and *λ_d_* for fixed values of *ω*.

We first study the performance of the NG, gravity, and radiation null models on temporally stable uniform population benchmarks (see Fig. 4) with parameter values *N* = 50, *l =* 10, and *m =* 10 layers using *γ* ∈ {0.5, 0.75, 1, 1.25, 1.5} and *ω* ∈ {10^-3^, 0.1, 0.25, 0.5, 0.75, 1}. We expect that for larger *ω* values the weight of the interlayer edges outweighs the intralayer edges, leading to each node being assigned to the same community as its copies in other layers. However, for the temporally stable benchmarks we did not observe this effect; here, we only show figures for *ω* = 0.1, as different values of *ω* give very little difference in results (in some plots nearly unnoticeable).

**FIG. 4.**
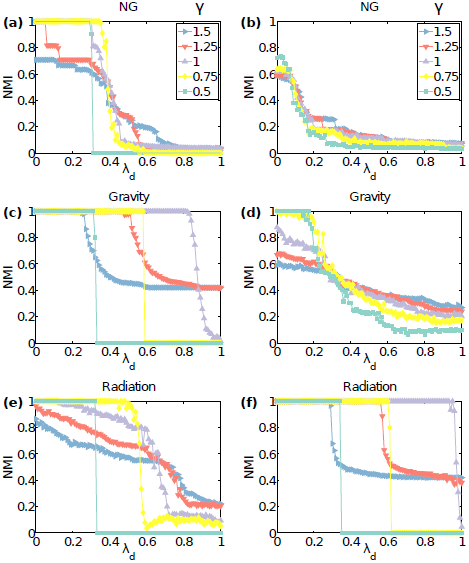
NMI between algorithmically detected and planted community structures in uniform population (*n_i_* = 100 for all *i*) multilayer temporally stable spatial benchmarks with *N* = 50, *l* = 10, *m* = 10, and *μ* = 100 for *ω* = 0.1 and various values of *γ* (colored curves) as a function of *λ_d_* for (left) the distance benchmark and (right) the flux benchmark. We detect communities by optimizing modularity using the (top) NG, (middle) gravity, and (bottom) radiation null models.

We also experimented with “random population” benchmarks (see Appendix E) and smaller and larger values of *ω*. Our results on multilayer benchmarks follow our findings from static benchmarks. Once again, we find that the choice of *γ* has a large influence on the quality of the algorithmic partitions, and (as with our findings for static benchmarks) *γ* = 1 seems to yield the best performance (i.e., the highest NMI scores) in most cases, except the near-singleton regime, where *γ* = 1.5 outperforms it slightly.

We now examine the NMI between algorithmic versus planted partitions on temporally stable multilayer benchmarks while varying *ω* and *λ_d_* for fixed *γ* = 1. As we show in Fig. 5, we find that the value of *ω* usually has little effect on our ability to detect the planted communities via modularity maximization on benchmarks with a temporally stable community structure. This suggests that the small interlayer variation due to the independent creation of layers is not enough to observe the influence of *ω* on community detection.

**FIG. 5.**
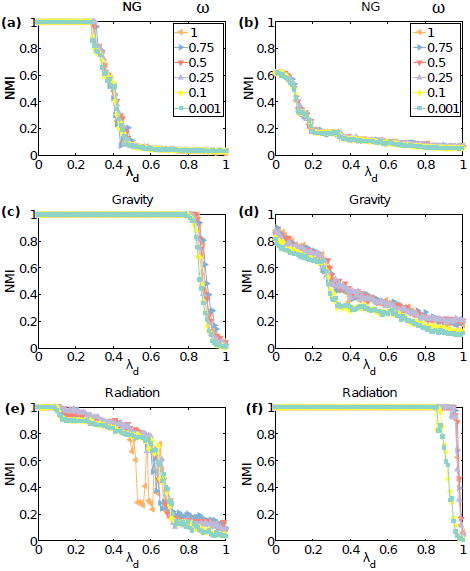
NMI between algorithmically detected and planted community structures in uniform population (*n_i_* = 100 for all *i*) multilayer temporally stable spatial benchmarks with *N* = 50, *l* = 10, *m* = 10, and *μ =* 100 for *γ =* 1 and different values of interlayer edge weights *ω* (colored curves) as a function of *λ_d_* for (left) the distance benchmark and (right) the flux benchmark. We detect communities by optimizing modularity using the(top) NG, (middle) gravity, and (bottom) radiation null models.

We then study the performance of the three null models on temporally evolving uniform population benchmarks (see Fig. 6) with parameter values of *N* = 50 nodes, a lattice parameter of *l* = 10, a fraction *p* = 0.4 of nodes that change community over the whole timeline, and *m* = 10 layers. We show results for *γ* ∈ {0.5, 0.75, 1, 1.25, 1.5} for *ω* = 0.1 and for *ω* ∈ {10^-3^, 0.1, 0.25, 0.5, 0.75, 1} for *γ* = 1. Compare Fig. 6 to the left panels of Figs. 4 and 5. We observe on temporally evolving benchmarks that varying *ω* makes a difference, where the structures for *ω* 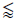 0.1 for the gravity null model and *ω* 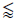 0.5 for the radiation null model are the most similar to the planted partitions. This is in accordance with our expectation that algorithmically detected community structure becomes overly biased towards connecting copies of nodes across slices above a critical *ω* value (which depends on network structure).

**FIG. 6.**
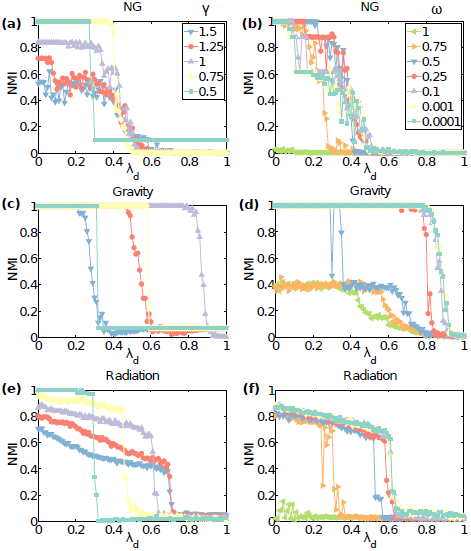
NMI between algorithmically detected and planted community structures in uniform population (*n_i_* = 100 for all i) multilayer temporally evolving spatial distance benchmarks with *N* = 50, *l* = 10, *m* = 10, and *μ* = 100 for (left) *ω* = 0.1 and different values of the resolution parameter *γ* (colored curves) and (right) *γ* = 1 and different values of the interlayer weights *omega* (colored curves) as a function of *λ_d_*. We detect communities by optimizing modularity using the (top) NG, (middle) gravity, and (bottom) radiation null models.

We also perform a “province-level” community detection on the multilayer benchmarks in which we seek assignments of nodes (regardless of what layer they are in) to communities and compare the results to benchmark networks with planted community structure. This is analogous to trying to detect community structure in disease data that persists over time — e.g., to seek the influence of climate on disease patterns. This is easiest to apply to temporally stable multilayer networks.

For temporally stable multilayer benchmarks (see the discussion in Appendix F), we successfully detect the underlying communities, and we obtain similar performance results as with the multilayer communities that we discussed above.

Our results on synthetic benchmark networks suggest that using a spatial null model on a spatial network does not necessarily assure a better result for community detection. The quality of results with different null models depends strongly on the data and the choice of parameter values. For example, incorporating population information into a null model in a situation in which the population is not influencing connectivity structure might cause community detection to yield spurious communities (as we discussed in the context of random population benchmarks).

The level of influence of different node properties or events (such as disease flux on edge placement) and the extent of mixing between communities is often unknown for networks that are constructed from real data. For such networks, we recommend to try both spatial and non-spatial null models over a wide parameter range and to study the results carefully in light of any other known information about the network. In Section IV, we will present an example of using such a procedure to study the community structure of correlation networks that are created from time series of dengue fever cases.

## IV. Application to Disease Data

In this section, we assess the performance of the NG, gravity, radiation, and correlation^2^ null models on multilayer correlation networks that we construct from disease incidence data that describe the spatiotemporal spread of dengue fever in Peru from 1994 to 2008.

Disease dynamics are strongly influenced by space, as the distance between regions affects the migrations of both humans and mosquito host [53]. They are also affected by climate due to the temperature dependence of the mosquito life cycle [64], and different regions of Peru have different climates. Therefore it is important to examine and evaluate the performance of different spatial null models when examining communities in networks that are constructed from disease data.

### A. The Disease and the Data

Dengue is a human viral infection that is prevalent in most tropical countries and is carried primarily by the *Aedes aegypti* mosquito [65]. The dengue virus has four strains (DENV-1-DENV-4). Infection with one strain is usually mild or asymptomatic, and it gives immunity to that strain, but subsequent infection with another strain is usually associated with more severe disease [65].

Although dengue was considered to be nearing extinction in the 1970s, increased human mobility and mosquito abundance have led to its resurgence in many countries — often as recurrent epidemics with an increasing number of cases and severity of disease. Dengue is a rising threat in tropical and subtropical climates due to the introduction of new virus strains into many countries and to the rise in mosquito prevalence from the cancellation of mosquito eradication programs [66]. It is currently the most prevalent vector-borne disease in the Americas [66, 67].

Peru is located on the Pacific coast of South America. Its population of about 29 million people is distributed heterogeneously throughout the country. The majority live in the western coastal plain, and there are much smaller population densities in the Andes mountains in the center and the Amazon jungle in the east. The climate varies from dry along the coast to tropical in the Amazon and cold in the Andes. Such heterogeneities influence dengue transmission [68]. For example, temperature [69] and rain [70] affect the life cycle of the main dengue vector *Ae. aegypti*, and temperature affects its role in disease transmission [71–73]. The jungle forms a reservoir of endemic disease; from there, the disease occasionally spreads across the country in an epidemic [64]. Additionally, as *Ae. aegypti* typically travels short distances [74], human mobility can contribute significantly to the heterogeneous transmission patterns of dengue at all spatial scales [57].

Our dengue data set consists of 15 years of weekly measurements of the number of disease cases across 79 provinces of Peru collected by the Peruvian Ministry of Health [75] between 1994 and 2008. These data have previously been analyzed by Chowell et al. to study the relationship between the basic reproductive number, disease attack rate, and climate and populations of provinces [64].

Until 1995, the DENV-1 strain dominated Peru; it mostly caused rare and isolated outbreaks [67]. The DENV-2 strain was first observed in 1995–1996, when it caused an isolated large epidemic [76]. DENV-3 and 4 entered Peru in 1999 and led to a countrywide epidemic in 2000–2001 [77], and there was subsequent sustained yearly transmission [67]. The data contains a total of 86,631 dengue cases; most of them are in jungle and coastal provinces (47% and 49%, respectively), and only 4% of the cases occur in the mountains. The disease is present in 79 of the 195 provinces.

In this paper, we use the definition of “epidemic” from the US Agency for International Development (USAID): an *ePidemiC* occurs when the disease count is two standard deviations above the baseline (i.e., mean) [78]. When stating countrywide epidemics, we apply this definition when considering all nodes. When stating local epidemics, we apply this definition to individual provinces (though one could also consider particular sets of provinces).

### B. Network Construction

Our data set *D* consists of *N* = 79 time series of weekly disease counts {*D*_1_, *D*_2_, …, *D_N_*} over *T* = 780 weeks. The quantity *D_i_*(*t*) denotes the number of disease cases in province *i* at time *t*. (See Fig. 7 for a plot of the number of cases versus time.) We create networks from this data by calculating the Pearson correlation coefficient between each pair of time series.^3^

**FIG. 7.**
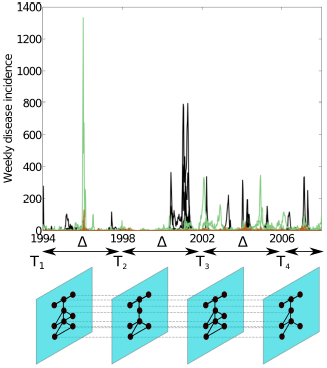
Construction of multislice correlation networks from disease time-series data. The top panel shows the dengue fever time series for the 79 provinces of Peru. We color the provinces by climate: coastal provinces are in black, mountainous provinces are in brown, and jungle provinces are in green. Observe the large epidemics in 1996 (focused in the jungle Utcubamba province) and 2000–2001 (countrywide, but primarily on the northern coast), and the recurrent post-2001 epidemics (which affect various jungle and coastal provinces). The bottom panel shows an example of the multislice network construction for 9 nodes with *τ* = {1, 209, 417, 625} and Δ = 208. (The time points correspond to 1/1/1994, 27/12/1997, 22/12/2001, and 17/12/2005). The nodes represent provinces and each intralayer edge weight is given by a Pearson correlation between a pair of single-province time series in a given time window. One set of correlations gives one temporal layer, and we connect copies of each node in neighboring layers using interlayer edges of uniform weight *ω* ∈ [0, ∞] (dashed lines). The case *ω* = 0 yields a set of static networks. (All other aspects of our network construction are the same.)

We seek to study the temporal evolution of the correlations by constructing separate networks for different time windows — we either construct a set of static networks or a multislice network. To create these networks, we divide each of the time series into *ω* time windows by explicitly defining a list of the starting points *τ* = {*τ*_1_, *τ*_2_, …, *τ_m_*} for each time window and the time window width Δ = *τ*_t+1_ - *τ_t_*. In the present paper, we use *τ*_1_ = 1 unless we state otherwise.

The starting point *τ_t_* and window width Δ define a time window that we use to select on portion of the disease time series. For example, for the time series of disease cases in province *i*, the time-series portion *E_i_* = {*D_i_*(*τ_t_*), *D_i_*(*τ_t_*+1),…, *D_i_*(*τ_t_*+Δ)} represents the numbers of disease cases in province *i* at times *τ_t_, τ_t_* + 1, …, *τ_t_* + Δ. By considering all provinces, one can use such time series either to construct a set of static networks or a multislice network.

For a static network, we define a set of *N* nodes {1, 2, …, *N*}, where node *i* corresponds to province *i*. The edge weight

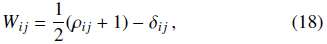

where the Kronecker delta *δ_ij_* removes self-edges, represents the similarity between the time series *E_i_* and *E_j_*. The quantity *ρ_i__j_* is the Pearson correlation coefficient between the disease time series for provinces *i* and *j*. That is,

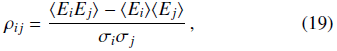

where <•> indicates averaging over the time window under consideration, and *σ_i_* is the standard deviation of *E_i_.* Our construction yields a fully connected (or almost fully connected) network *W* with elements *W_ij_* ∈ [0, 1]. When studying static networks, we use *τ* = {1, 2, …, *T* - Δ } to form a set of *T* - Δ overlapping static networks.

To construct a multislice network, we use the times 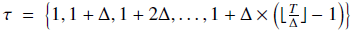 to create nonoverlapping time windows. The intralayer edge weights are

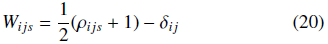

for each layer *s*. We connect each node *i* in the *r*th time window to copies of itself in an adjacent time window *s* using interlayer edges of uniform weight *C_isr_ = ω* ∈ [0, ∞]. This yields a weighted multislice correlation network. The case *ω* = 0 in the multislice network corresponds to a sequence of static networks. See Fig. 7 for a schematic that shows the construction of a multislice network.

Similar constructions of (both static and multislice) networks from time series have been employed for systems such as functional brain networks [27, 79], currency exchange-rate networks [22], and political voting networks [24, 80, 81].

Many features, such as the number of layers and the mean and variance of the Pearson correlation values, depend on the parameters that we use in constructing our networks. For example, it is important to consider the choice of the time window size Δ. There is a trade-off between having many layers to obtain a good temporal resolution of events and ensuring that we construct each layer using enough time points to be confident of the statistical significance of the similarity values in the adjacency-tensor layer [79]. Larger values of Δ yield smaller variations in mean correlation across the years and lessen the effects of small, regional epidemics on the number of cases and on the correlation between disease profiles in different provinces.

Therefore, we want to use a sufficiently large value of Δ so that we can examine long-term, repetitive disease patterns. However, choosing a value of Δ that is too large risks over-smoothing the data and losing important information.^4^ In the present study, we investigate long-term patterns of disease spread. Unless we state otherwise, we use Δ = 52 for the (overlapping) static networks in order to observe yearly disease patterns. However, we use Δ = 26 for multilayer networks to ensure that we have enough layers to study the temporal evolution of disease patterns and that the yearly epidemic peak is in only one of the two layers that cover that year. Because our disease data starts on 1 January and the yearly disease peaks usually appear in the first 20 weeks of a given year, all but one of the observed peaks only span one layer.

### C. Community Structure in Disease-Correlation Networks

It is well-known that geographical distance has an important influence on disease spread [57, 83, 84]. Additionally, climate exerts a significant influence on dengue, and it is also necessary to consider Peru’s particular topography (as its mountains form a barrier to disease spread) [64, 67]. Therefore, we expect the community structure in the disease-spread networks to be strongly geographical. We also expect to observe large changes in community structure at certain time points — such as when the introduction of the new disease strains around 1999 led to large epidemics and the onset of yearly countrywide epidemics [67]. In this section, we explore the similarity of algorithmically obtained community structures to spatial and temporal groupings of nodes across a range of parameter values.

To compare the algorithmic partitions of the correlation networks versus manual partitions, we use the *z*-score of the Rand coefficient [61]. The Rand coefficient is

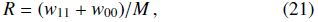

where *w*_11_ is the number of node pairs that are in the same community in both partitions, *w*_00_ is the number of node pairs that are in different communities in both partitions, and *M* is the total number of node pairs.

We use *z*-Rand scores instead of NMI because the former measure is good at detecting similarities in coarse structure [10, 61] but is less sensitive to minor changes such as one node changing community assignment. For the disease data, we do not possess ground-truth partitions as we did for our synthetic benchmark examples, so we seek to evaluate broad organizational similarities in the algorithmic and manual partitions rather than attempting to conduct a fine-grained evaluation of community structure versus a planted partition. We thereby aim to inform our understanding of the general structural influences on the spatiotemporal patterns of disease spread. One can also examine measures of spatial autocorrelation (e.g., Moran’s *I*) [85].

To examine the spatial community structures in the static and multilayer networks, we compare the results of the partitions that we obtain algorithmically to manual partitions using *z-*Rand scores. In the “climate partitions”, we group nodes according to the topography of their associated provinces — jungle, coastal, and mountainous provinces — and then subsequently divide the coastal and mountainous communities into northern, central, and southern provinces [see Fig. 8(a,b)]. We use the detailed climate partition for the subsequent study. In the 19-community “administrative partition”, we assign each node to its associated administrative region [see Fig. 8(c)]. We compare each of the 728 static networks against these two manual partitions to study the spatial element of the data. For the multilayer networks, we compare the algorithmic partition versus a manual partition by taking the same manual partition of nodes for all slices.

**FIG. 8.**
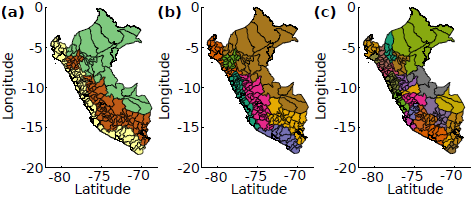
Visualization of the three different topographical partitions of Peru’s provinces on a map. (Left) Broad climate partition into coast (yellow), mountains (brown), and jungle (green); (center) the further division of coast and mountains into northern coast, central coast, southern coast, northern mountains, central mountains, and southern mountains; and (right) the administrative partition of Peru.

We use the term “spatial partitions” to describe partitions that yield high *z-*Rand scores in comparison to the climate or administrative manual partitions. For multilayer networks, we also compare the algorithmic partitions to partitions that contain a planted temporal change in community structure. For these comparisons, we group the multilayer nodes into ones that occur before or after a “critical” time *t_c_*, and we use the term “temporal partitions” to describe partitions that yield high *z-*Rand scores in this comparison. We test all the times 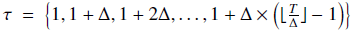 that we use to create the multilayer network, and we report the time with the highest *z-*Rand score as the critical time *t_c_*.

#### 1. Community Structure Using the NG Null Model

Before looking at multilayer networks, we first study the community structures of the 728 overlapping static networks formed by taking *τ* = {1, 2, …, 728} and using Δ = 52. We select the networks for which the algorithmic partitions score the highest against manual spatial partitions of the network for further study.

The community structures that we obtain from maximizing modularity have a strong spatial element, as suggested by the high *z*-Rand scores when compared to topographical partitions. As one can see in Fig. 9(a), which shows a compact box plot of the *z*-Rand scores versus climate partitions for resolution parameter values of *γ* ∈ 0.1, 0.2,…, 3 (each box) across the 728 networks covering the data set (the horizontal axis), the spatial element is especially evident after the year 2000.

As one can see from a plot of number of epidemic cases over time (see Fig. 7), this transition seems to occur around the time of the largest countrywide epidemic in the data, and the subsequent period includes recurring yearly epidemics that have been linked to climatic patterns in prior studies [67]. There are two periods of significantly spatial partitions: one corresponds to the 2000–2001 epidemic, and the second occurs in 2002–2004, which contains the spatial partition with the highest *z-*Rand score against climate [see Fig. 9(b)]. Note that the topographical *z-*Rand scores decrease after 2004 despite the continuing yearly dengue epidemics.

By plotting the partitions that have the highest *z-*scores with respect to the manual climate and administrative partitions on a map of Peru in Figs. 9(b,c), we observe that the statistically significant similarity of the algorithmic partition to these partitions is difficult to discern by eye.

This is largely due to the large number of singleton communities (38 of 47 communities for the highest-scoring climate partition, and 60 of 64 communities for the administrative partition). One possible cause for the large number of singleton communities is that epidemics have only occurred every few years on a province scale (so many provinces thus have somewhat independent disease histories), although there are sustained transmissions and yearly epidemics on a countrywide scale. This might be due to the independent development of immunity of the four strains of dengue, which could cause the populations of the different provinces to be susceptible to different strains of the disease, which could in turn lead to epidemics that are dominated by one serotype each year (as has occurred with dengue in Thailand) [86]. In this scenario, one would not expect all epidemics to reach every province (which could, in turn, lead to singleton communities).

The jungle nodes form the largest communities in these spatial partitions, and these contribute the most to the high spatial scores. In the time periods covered by the two static networks corresponding to the highest-scoring administrative and climate partitions, there was a dengue epidemic in some jungle provinces at the time corresponding to the static network (see Fig. 7). Indeed, seven of the twelve jungle nodes in the May 2003 partition (six of which are located in close proximity to each other) experienced a dengue epidemic for six weeks in the year corresponding to that network. It is possible that the proximity is driving the high synchronization in epidemic spread between these provinces.

Let us now consider community structure in the multilayer disease network with nonoverlapping layers that we construct using the time points *τ* = {1, 27, …, 755} and using Δ = 26. To find interesting parameter regimes, we compare the algorithmically computed community structure of the dengue fever multilayer disease-correlation network to manual partitions across a range of *ω* and *γ* parameter values between 0 and 3 (see Fig. 10). For *γ* 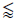 1, all nodes are in one community. For *γ* ∈ [1, 1.5] and *ω 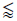* 1, the community structure scores the highest versus the temporal partition [see Fig. 10(c)]. The community structures for *γ* ∈ [1, 1.5] (where the endpoints of this interval are approximate) and *ω 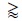* 1 exhibit a mixture of spatial and temporal features. For *γ* 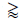 2, the multilayer community structure has high *z-*Rand scores (they are larger than 100) in comparison to climate partitions. This results from the large value of interlayer coupling between corresponding nodes in consecutive layers.

**FIG. 9.**
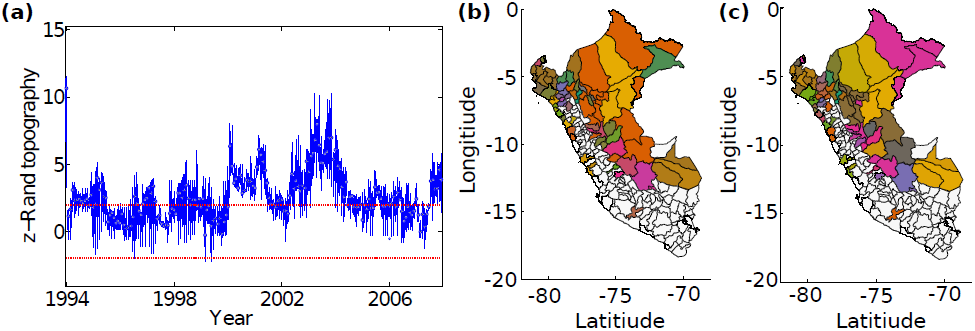
Properties of algorithmic community structure, which we obtained by maximizing modularity using the NG null model for the dengue fever static correlation networks with window size Δ = 52. (a) A box plot of the *z-*Rand scores versus the detailed climate partition at different *γ* values (*γ* ∈ 0.1, 0.2, …, 3), for the 728 static networks covering the whole time period (horizontal axis). The red lines show *z-*Rand scores of ±1.96 for guidance. (b) Community structure with the highest *z-*Rand score when compared to the climate partition. The resolution-parameter value is *γ* = 0.5, the layer is 492 (which occurs in May 2003), the *z-*Rand score is 9.65, and we show the largest community in orange. (c) Community structure with the highest *z-*Rand score when compared to the administrative partition. The resolution-parameter value is *γ* = 0.5, the layer is 1 (which occurs in January 1994), the *z-*Rand score is 11.53, and we show the largest community in brown. Our visualization in panels (b,c) uses a map of Peru in which we color provinces according to their community assignment. White provinces are ones in which our data does not include any reported cases of dengue fever in the indicated time window.

When studying the qualitative features of the partitions for *γ* ∈ [1, 1.5] (where the endpoints of this interval are approximate) and *ω* 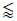 1, we observe that community detection repeatedly finds that January 2002 is the critical time *t_c_* (i.e., the strongest change point in temporal community structure). This finding suggests that a strong shift in the patterns of disease correlations occurred around this time. Indeed, Peru experienced a large countrywide dengue epidemic in 2000–2001, and this period also marks the onset of new yearly epidemic dynamics [67]. Thus, our method recovers the most important biological event in this data set in addition to providing additional information about spatial influences on disease spread. We also observe several other time when new communities are born: June 1999 (the first large epidemic after 1996), January 2004, and January 2007. We do not know the biological significance of the latter two dates. Notably, in this parameter regime, our community structure does not identify the large epidemic in the jungle Utcubamba province in 1996 (see Fig. 7), which is the other large event in this data set.

The community structure that we detect depends heavily on parameter values. In many parameter regimes — especially when *γ* 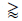 1 and *ω* 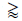 0.5 — communities appear to be predominantly spatial, and we find high *z-*Rand scores when compared to the climate and administrative partitions [see Fig. 10(b)]. The high influence of spatial proximity on the community structure is unsurprising, as spatial distance is an important influence on disease spread [57, 84]. Previous studies have also noted that the community structure of spatial networks obtained by maximizing modularity using the NG null model tends to be strongly influenced by geographical factors [20, 87, 88]. If there are other interactions that shape the dengue fever correlation network, they might be obscured by the strong influence of spatial proximity. However, such interactions might be revealed by using a spatial null model that incorporates the expected effect of space on interactions. We pursue this idea in Section IV C 2.

**FIG. 10.**
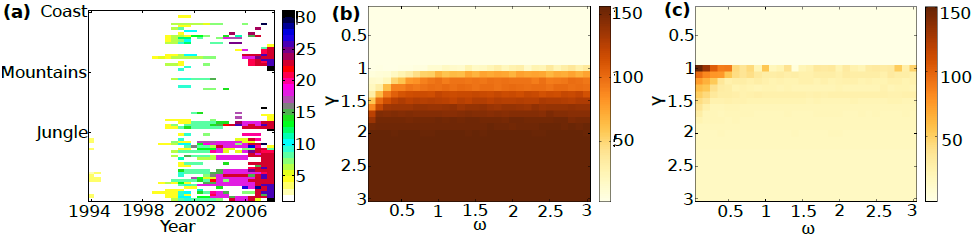
Algorithmic partitions, which we obtain by maximizing modularity using the NG null model, of the dengue fever multilayer disease-correlation network that we construct using Δ = 26. (a) An example of a consensus persistent community structure that we obtain for a resolution-parameter value *γ* = 1 and interlayer coupling values of *ω* ∈ [0.1, 3]. Layer start times *τ* are plotted on the horizontal axis, and nodes are on the vertical axis. Node community membership is indicated by color. We observe several times when communities die and new ones are born. (b,c) Results of varying the parameters *γ* and *ω*. We show the *z*-Rand scores for similarity to (b) “spatial” partitions by climate and (c) temporal partitions before and after a critical time *t_c_*. (For this figure and for each set of parameter values, we select the highest scoring *t_c_*; in the majority of cases, *t_c_* corresponds to January 2002.)

#### 2. Community Structure Using Spatial Null Models

We apply spatial null models to the dengue fever correlation networks. We obtained province locations from the Geonames.org website [89], and we obtained their populations from the Peruvian Instituto Nacional de Estadística e Informática (INEI) [75]. We were only able to obtain the 1994 and 2007 populations; due to the limited range of data and the several changes in Peruvian administrative structures between the two times, we only use the 2007 populations.

The maximum inter-province distance is about 1300 km. We report numerical experiments using a bin size of 100 km after testing the spatial deterrence for several other sizes (ranging between 50 and 500 km) in the same manner as in Ref. [20]: that is, we study the shape of the deterrence function [see Eq. 5 and the nearby discussion] with changing distance across bin sizes, and we then examine the community structures that we obtain using different bin sizes. We find that bin sizes have an effect on the shape of the deterrent function (with lower sizes giving smoother results), but all of the bin sizes that we tested produced very similar partitions for both the gravity and radiation spatial null models.

Recall from Section IV A that only 79 of the 195 provinces had reported cases of dengue fever in our data, so we use the location and population data only for those provinces.

We first study the community structure on static disease-correlation networks using the gravity and radiation null models. Both null models seem to most remove the spatial element of the community structures (including any temporal variation in the spatial correlations), as indicated by a lack of variation the spatial *z*-Rand scores (not shown). For both the gravity and radiation null models, we observe high similarity between layers for a variety of values of the resolution parameter *γ* [see Fig. 11(a)]. These structures contain one dominant community with over 70 nodes along with several singleton communities [see Fig. 11(b)]. By examining the partitions directly using a map of Peru, we see that the singleton communities tend to consist of the highest-populated nodes. For example, we obtain the largest *z*-Rand score with respect to a climate partition for the gravity model when the Lima province, which contains 41% of the country’s population, is a singleton community [see Fig. 11(c)].

**FIG. 11.**
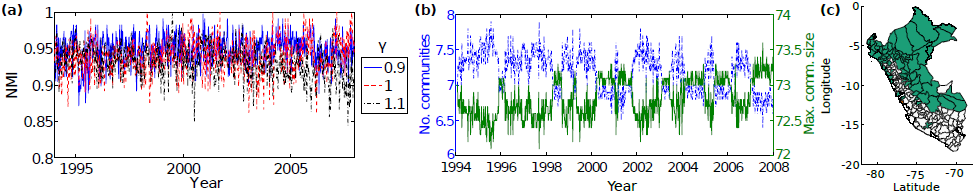
Properties of the algorithmic community structure, which we detected by maximizing modularity using the gravity null model, of the dengue fever static correlation networks that we construct using a time window of Δ =52. (a) NMI between adjacent layers for *γ* ∈ {0.9, 1, 1.1}. (b) Maximum community size (green solid curve) and number of communities (blue dashed curve) for *γ* = 1. (c) Community structure scoring the highest *z*-Rand score versus climate among the dengue fever static correlation networks that we construct using Δ = 52. (The resolution-parameter value is *γ* = 1, the layer is 45, and the *z*-Rand score is 2.60.) We show these structures on a map of Peru, and we color provinces according to their community assignment. White provinces are ones in which our data does not include any reported cases of dengue fever in the indicated time window. Observe the single giant community that contains almost all of the nodes except the Lima province (which is a singleton with 41% if the population).

We also examine the spatial null models for multilayer correlation networks. The community structures again exhibit one large community containing the majority of multilayer nodes [see Fig. 12(a,b)], and several multilayer nodes corresponding to provinces with highest populations form singleton communities across time. This situation occurs for all of the tested parameter values. Additionally, we do not observe any clear pattern in the *z*-Rand scores as we change *γ* and *ω*.

**FIG. 12.**
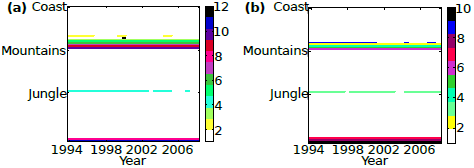
Consensus community structure, which we obtain by maximizing modularity using (a) the gravity null model and (b) the radiation null model, of the dengue fever multilayer disease-correlation network that we constructed using a time window of Δ = 26. We use a resolution-parameter value of *γ* = 1 and consider *ω* ∈ [0.1, 3]. Layer start times *τ* are plotted on the horizontal axis, and nodes are on the vertical axis. Node community membership is indicated by color.

Our findings suggest that the addition of space into a null model for modularity optimization might remove the majority of the variation in the correlation structure of the dengue fever correlation networks, such that the influence of population size could be the only major factor that remains. This could relate to the concept that a minimum population size is required for sustained disease transmission; it has been estimated that this size is between 10,000 and 500,000 for dengue [64, 90]. There are only 5 provinces with populations over 500,000, and these provinces are often assigned to singleton communities when we use a spatial null model. This suggests that they have different disease patterns from the other provinces.

### D. Community Detection Using a Correlation Null Model

Recently, MacMahon et al. [18] proposed a new null model that they designed specifically for modularity maximization for networks that are constructed based on the pairwise Pearson correlations between time series. They used ideas from random matrix theory (RMT) [91] to generate a null model that represents the “random” component of a correlation matrix and can take into account the single most strongly influencing factor on the correlation structure. In the context of financial systems, which was the focal example of Ref. [18], this factor is often called a “market mode”. Given that we often found a single large community when we used spatial null models, it is interesting to see what results we obtain using such a correlation null model.

To use a correlation null model, we need to construct our network directly from pairwise correlations without subsequently shifting them to [0, 1] and removing self-edges. We construct networks by selecting time windows and calculating Pearson correlations in the same manner as in Section IV B, but the here edge weights are left as raw correlations: *C_i__j_ = ρ_i__j_* (Eq. 19).

Because of the special structure of correlation matrices, modularity using the standard NG null model assigns importance to pairs of nodes *i* and *j* whose Pearson correlation is larger than the product of the correlations of each node with the time series of the total number of disease cases in the country over the chosen time window: *E*_tot_, where *E*_tot_(*t*) = 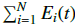.

By contrast, the correlation null model that we adopt from Ref. [18] uses ideas from RMT to detect communities of nodes that are more connected than expected under the null hypothesis that all time series are independent of each other.

For a given correlation matrix constructed from *N* time series that each have length *T* (with *T* /*N* > 1), one posits based on RMT that any eigenvalues that are smaller than the eigenvalue 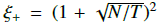 are due to noise. Here, *ξ*_+_ is the maximum eigenvalue predicted for a correlation matrix that is constructed from the same number of entirely random time series.

Additionally, for many empirical correlation matrices (including ours, as we show in Table II), the largest eigenvalue ξ_m_is much larger than the others, and its corresponding eigenvector has all positive signs [18]. In this situation, there is a common factor, which is called the “market mode” in financial applications, that influences all of the time series [92].

**TABLE II.** Characteristics of disease-correlation networks that we construct using different time window widths Δ, including the mean *ξ_m_* and the mean number of eigenvalues *ξ* that are larger than *ξ_+_* and are thus deemed to correspond to non-random elements of the matrix. The networks below the horizontal line satisfy *T*/*N* > 1 and always have more than one eigenvalue ξ per layer that satisfies *ξ* > *ξ_+_*.

| N | $\Delta$ | No. Layers | $\xi_+$ | $\xi_m$ | No. $\xi > \xi_+$ per layer |
| --- | --- | --- | --- | --- | --- |
| 79 | 26 | 754 | 4.36 | 5.99 | 1.01 |
| 79 | 52 | 728 | 3.99 | 5.84 | 1.07 |
| 79 | 80 | 700 | 3.975 | 6.24 | 1.30 |
| 79 | 104 | 676 | 3.50 | 5.61 | 2.13 |
| 79 | 130 | 650 | 2.47 | 5.41 | 2.19 |
| 79 | 150 | 630 | 2.28 | 5.26 | 2.31 |

We can thus decompose our correlation matrix *C* as follows: *C* = *C*^(^*^r^*^)^ + *C*^(^*^g^*^)^ + *C*^(^*^m^*^)^, where *C*^(^*^r^*^)^ is the “random” component of the matrix, *C*^(^*^m^*^)^ is the “market mode”, and the “group mode” *C*^(^*^g^*^)^ is embodies the meaningful correlations between time series. We write 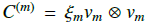 and 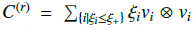, where *ξ_i_* and *ν_i_* are an eigenvalue and its corresponding eigenvector, *ν_m_* ⊗ *ν_m_* is the outer product of the two vectors (a special case of the Kronecker product for matrices), and *ξ_m_* is the maximum observed eigenvalue in the correlation matrix *C*. We can construct a correlation null model either by removing both the “random” component of the matrix and the influence of the “market mode” (i.e., by using the null model *P*^corr^ = *C*^(^*^r^*^)^ +*C*^(^*^m^*^)^) or by only removing the random component (i.e., by using the null model *P*^corr^ = *C*^(^*^r^*^)^).

As a necessary preliminary calculation, we examine the maximum eigenvalue *ξ_m_* and the relationship between the values of *T* and *N* for the dengue fever correlation networks with time windows of sizes Δ = 26 and Δ = 52. To satisfy the *T*/*N* > 1 requirement to applying the RMT approach of Ref. [18], we require Δ ≥ 80. However, for Δ = 80 as well as for Δ ∈ {26, 52} not all layers contain more than one eigenvalue *ξ* that satisfies *ξ* > *ξ_+_* (see Table II), and most of the correlation matrix is classified as noise. To avoid this, we choose to consider larger Δ ∈ {104, 130, 150}. For subsequent calculations, we use Δ = 104 unless stated otherwise. In Appendix G, we compare these results to our results for Δ ∈ {26, 52, 130, 150}.

Although the maximum eigenvalues that we discussed above are much larger than the other eigenvalues and every component of the associated eigenvector is positive, the eigenvector does not affect all nodes to the same extent. The above construction thus yields a non-uniform null model for our data in practice, so we are unable to identify the analog of a market mode. We thus do not incorporate such a mode into the null models that we employ for community detection. We use the correlation null model

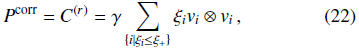

where *γ* is the resolution parameter. For the multilayer setting, we write

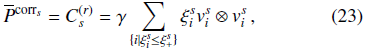

where 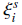 and 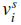 are an eigenvalue and its corresponding eigenvector for layer *s*.

We test the performance of this correlation null model on correlation networks that we construct from dengue fever time series with Δ = 104. In most of the static networks, the community structures appear to be affected by spatial proximity — especially for post-2000 networks, as illustrated by the high *z*-Rand scores versus the climate partition (particularly in 2000–2001, 2002–2004, 2005–2006). See Fig. 13(a). These high *z*-Rand scores result from (1) the classification of the majority of jungle provinces into one community and (2) the existence of a community that contains many of the northern coastal provinces [see Figs. 13(b,c)]. We obtain similar results for Δ = 130 and Δ = 150, whereas the jungle provinces are split into two communities for Δ = 26 and Δ = 52. See Appendix G.

**FIG. 13.**
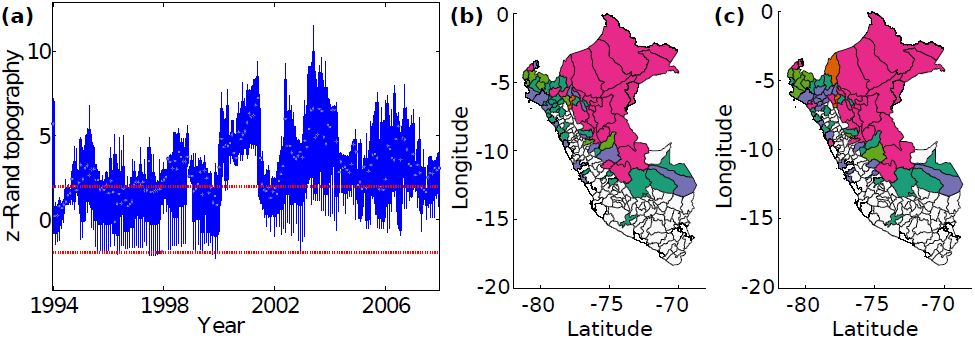
Algorithmic community structure, which we obtain by maximizing modularity using a correlation null model, for the static dengue fever correlation networks that we construct using Δ = 104. (a) A box plot of the *z*-Rand scores versus the detailed climate partition at different *γ* values (*γ* ∈ 0.1, 0.2, …, 3), for the static networks covering the whole time period (horizontal axis). The red lines show *z*-Rand scores of ±1.96 for guidance. In panels (b,c), we show partitions of the network with the highest *z*-Rand score based on (b) climate (*γ* = 1; layer 532, which corresponds to February 2004; and a *z*-Rand score of 8.9) and (c) administrative divisions (*γ* = 1; layer 492, which corresponds to May 2003; and a *z*-Rand score of 10.13). We color provinces according to their community membership on a map of Peru. White provinces are ones in which our data does not include any reported cases of dengue fever in the indicated time window.

We also perform community detection on multilayer networks using the correlation null model 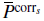 for (*γ, ω*) ∈ [0.1, 3] × [0.1, 3]. We obtain community structures with a mixture of temporal and spatial features. We calculate a consensus community structure for each given value of *γ* for the various values of *ω*. In Fig. 14(a), we show the best-looking persistent partition, which we obtain for *γ* = 1 and *ω* ∈ [0.1, 3]. This partition includes 15 communities. Although several communities coexist in each layer, the primary divisions appear to be largely temporal. For example, community 2 shrinks around 1998, and community 9 grows after 2005. The value of *t_c_* that yields the highest *z*-Rand score (against a temporal partition) is again January 2002 for this choice of (*γ, ω*), and this is also the case for the majority of (*γ, ω*) ∈ [0.1, 3] × [0.1, 3].

**FIG. 14.**
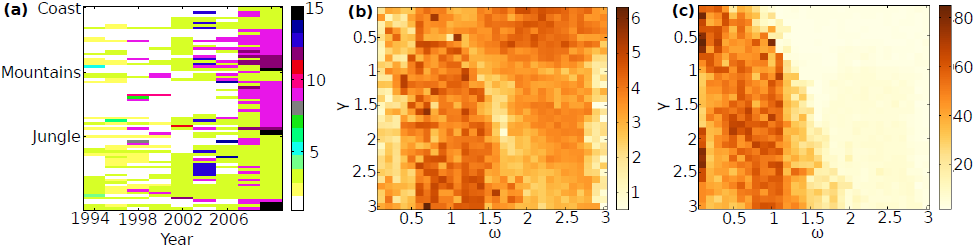
Algorithmic community structure, which we obtain by maximizing modularity using a correlation null model, of the dengue fever multilayer disease-correlation network that we construct using Δ = 104. (a) Consensus persistent community structure for *γ* = 1 for *ω* ∈ [0.1, 3]. (b,c) Results of varying *γ* and *ω*. We show the *z*-Rand scores for similarity to (b) “spatial” partitions by administrative region and (c) temporal partitions before and after a critical time *t_c_*. (For each parameter set, we select the highest scoring *t_c_*.)

In our sweep over the different values of *γ* and *ω*, we obtain relatively low climate *z*-Rand scores compared to what we obtained using NG null model (see Section IV C 1). We do not observe any clear patterns in the spatial *z*-Rand scores as we vary *γ* and *ω*, and we obtain relatively high temporal *z-*Rand scores (up to about 80) for *ω* 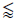 1 [see Fig. 14(b,c)]. For *ω* ∈ [1, 1.5] (where the endpoints of this interval are approximate), the larger values of *γ* values yield more temporal like partitions. For *ω* 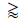 1.5, by contrast, the temporal *z-*Rand scores are well below 20.

#### 1. Province-level Multilayer Communities

We now examine the province-level information that we can glean from the data. The simplest approach is to construct a single static network from the entire length-*T* time series, but our multilayer approach allows us to aggregate data less severely. This, in turn, allows us to lose less information.

When we aggregate all time series to construct a single similarity network (i.e., we choose *τ* = 1 and Δ = 779), we find that the community structures that we obtain via modularity maximization with the spatial and correlation null models all consist of a single large homogenous community with up to three outlier nodes (see Fig. 26 in Appendix H). Only the NG null model is able to detect meaningful-looking communities, especially for *γ* = 1 and *γ* = 1.1 [see Fig. 15(a)]. For *γ* = 1, the we find two communities; the smaller one of them consists almost exclusively (15 of 18 nodes) of northern coastal provinces. This partition has *z*-Rand score versus climate of 7.3. For *γ* = 1.1, using the NG null model yields 28 communities, and many of them are small. The largest community corresponds exactly to the smaller community from *γ* = 1.

**FIG. 15.**
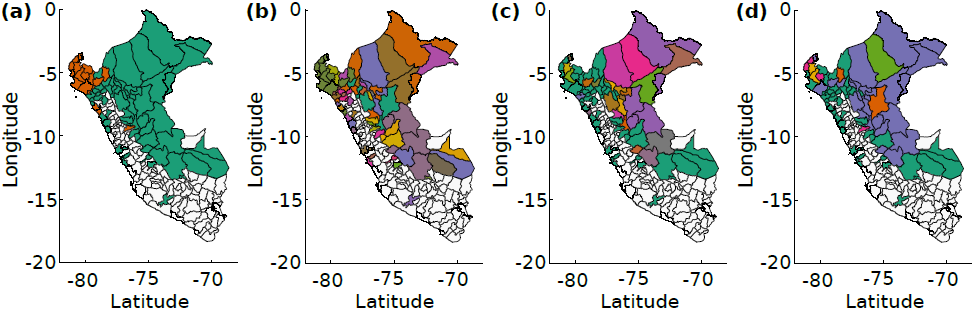
Province-level algorithmic community structure, which we obtain by maximizing modularity, for the static and multilayer dengue fever correlation networks. We color the provinces according to their community assignments. White provinces are ones in which our data does not include any reported cases of dengue fever in the indicated time window. (a) NG null model that is fully aggregated (i.e., *τ* = 1 and Δ = 779) with a resolution-parameter value of *γ* = 1. (b) NG null model that is fully aggregated with *γ* = 1.1. (c) NG null model in a multilayer network with province-level communities that we obtain from the multilayer network with a time window of width Δ = 26. (d) Correlation null model in a multilayer network with province-level communities that we obtain with a time window of width for Δ = 104.

Nodes grouped in the community of northern coastal provinces are the provinces of Peru that were most strongly involved in the 2000–2001 dengue epidemic; 15 nodes in this community experienced this epidemic, whereas only two other nodes experienced it.

The data aggregation over the whole time series results in the 2000–2001 epidemic dominating all other events in the time series. If we use the community structure of the temporally evolving multilayer network to create the province-level structure, we might be able to shed some more light on other interactions between provinces.

We then study the structure of province-level communities that we obtain from community detection using the uniform null model on an association matrix *A*^province^. As we discussed in Section II, we create this matrix by counting the number of multilayer nodes that are classified together in a consensus community detection on a multislice network. We consider the parameter values *γ* = 1 and *ω* ∈ [0.1, 3]).

Comparing the province-level communities that we obtain using the NG and correlation null models versus the broad topographical categories of coast, mountain, and jungle reveals the large-scale climatic influence on disease patterns. This is especially evident in the division into jungle and non-jungle provinces [see Fig. 15(c,d)]. We observe with the NG null model that the majority of mountainous and coastal nodes are classified together into one large community, in which jungle nodes are underrepresented (the p-value is 4.69 × 10^-6^ in a one-tailed Fisher exact test). As illustrated in Fig. 16(a), the jungle provinces are assigned to five singleton communities and six other small communities. With the correlation null model, we find that the coastal and mountainous provinces are again in one large community, in which the jungle nodes are underrepresented (the p-value is 2.48 × 10^-5^ in a one-tailed Fisher exact test), and the majority of jungle provinces are in a second large community in which jungle nodes are overrepresented (the p-value is 2.31 × 10^-6^ in a one-tailed Fisher exact test); see Fig. 16(b).

**FIG. 16.**
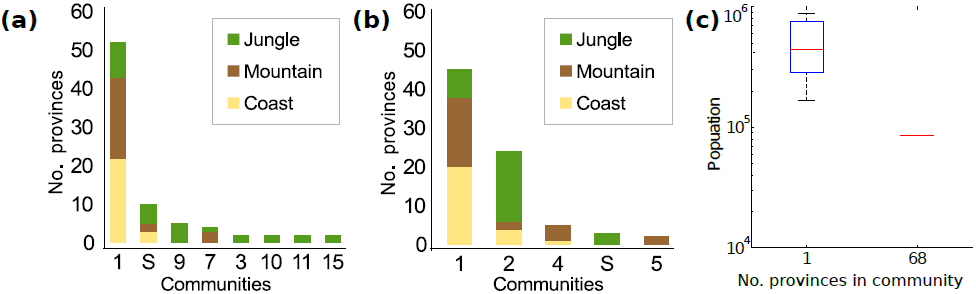
Membership of the consensus (across *ω* ∈ [0, 3]) province-level communities, which we computing by maximizing modularity, in multilayer dengue fever networks for *γ* = 1. In panels (a) and (b), we compare the climate composition of the communities using (a) the NG null model and (b) a correlation null model. We order communities according to their size, and the horizontal axis gives either the community number or “S” for a composite of the singleton communities. Most jungle nodes are combined into one community when using the correlation null model. (c) Box plot that indicates the population sizes of the communities that we find using the gravity null model. We group communities according to the number of provinces that they contain, and we observe that the singleton communities have populations that are much larger than the provinces that are assigned to the one large community.

The spatial null models that we study place almost all nodes in the same community. The only parameter that seems to influence community membership is population size, as the most populous nodes are outliers that form singleton communities at all examined values of *γ* and *ω* [see Fig. 16(c)].

## V. Conclusions

In conclusion, we examined time-dependent community structure, and we compared the use of different null models — including ones that incorporate spatial information — in the results of modularity maximization. We conducted our computational experiments using correlation networks constructed from spatiotemporal dengue fever incidence data in provinces of Peru (a system that is strongly influenced by spatial effects) and using novel synthetic benchmark spatial networks. We compared our results for the standard Newman-Girvan null model versus two null models that incorporate spatial information: a gravity null model [20] and a novel radiation null model. We also compared the NG null model on disease-correlation networks with a recently-developed correlation null model (and a multilayer generalization of it) that is designed specifically for studying correlation networks that are derived from time series [18].

Our results indicate that it is very important to incorporate problem-specific information such as spatial information into the null models for community detection. Our results also illustrate that there are many nuances to consider. That is, it is not simply a matter of incorporating spatial information in an arbitrary way but rather developing spatial null models that are motivated by application-appropriate generative models. For example, the NG null model performs better than the spatial null models (which both use population data) on the random population distance benchmark where populations vary but edge weight does not depend on them. However, when we remove the variation in population or modify the benchmark to include population in edge placement probabilities, we find that the gravity null model performs best (as expected).

Parameter choices can also be extremely important, as demonstrated by the large influence of bin size (when binning distances for the spatial null models) on community detection results, the failure to find meaningful communities with any of the null models at low edge densities, and the strong influence of resolution parameter *γ* on the results.

To summarize, one needs to consider seriously what variables that influence the connections in a system of interest that one wants to include in a null model, be careful about including spurious variables, and test how the results change for many parameter values.

Finally, not incorporating space at all can be more appropriate than incorporating it in a manner that is overly naive. (See, for example, our results on the random population benchmarks.)

In our consideration of dengue fever data, we observed for static networks that the NG and correlation null models find structures that are strongly spatial—especially after the onset of yearly epidemics in 2000. In our study, we observed that both yield partitions that include a large number of singleton communities and that spatial partitions are often dominated by large communities of neighboring jungle nodes that experience local epidemics during the time window.

On a multilayer network, maximizing NG modularity can result in either spatial or temporal partitions (depending on the parameter regime). Temporal partitions successfully find the most important time point in the history of the disease — namely, the introduction of a new disease strain that caused a large epidemic in 2000–2001 and a subsequent shift in disease patterns — and several other potentially interesting time points and periods of high spatial correlation.

When studying province-level connectivity, we illustrated that consensus province-level communities from an association matrix that we constructed from the multilayer network across time is a far preferable approach to complete data aggregation. For the aggregation into a static network, maximizing modularity using any of the test null models except the NG null model failed to detect any meaningful communities; the NG community structure corresponds to the large 2000–2001 epidemic. Aggregating networks results in loss of information that is desirable to study for meaningful patterns [3, 5].

When we constructed multilayer networks and computed consensus communities, the computed “spatial” multilayer partitions and province-level partitions highlight the importance of climate to the disease patterns of dengue, as the jungle provinces are placed into distinct communities from the mountainous and coastal provinces. This is sensible, as the yearly epidemic patterns tend (on average) to exhibit an earlier epidemic onset in the jungle [64, 67] and the jungle climate is rather distinct from the climate in coastal and mountainous provinces. The main climatic difference between jungle provinces and other provinces is temperature, and the influence of temperature on dengue transmission [68, 71, 72] and attack rate and persistence has been documented [64, 73]. The assignment of jungle provinces to communities is different for different null models. For the NG null model, they form small or singleton communities; for the correlation null model, they are grouped into one large community.

Which of these results is a more suitable grouping is debatable, as the jungle provinces tend to experience isolated epidemics; their disease time series have a lower mean Pearson correlation between themselves (0.0491) than the mean Pearson correlation between the jungle province time series and the entire data set (0.1116). However, grouping the jungle nodes into one community is also interesting, as it hints that the variables that influence jungle epidemics could be different than those in other climates. Chowell et al. [64] reported that the coastal and mountainous provinces exhibit more spatial heterogeneity of disease incidence than the jungle provinces, and population size appears to play a larger role in disease persistence in the jungle. Additionally, the jungle climate is more homogenous (especially in the north-south direction) than the other two climates.

When we attempt to remove the influence of space by using the gravity and radiation null models, we obtain one large community that contains all but the highest-population provinces (which are assigned to singleton communities). In contrast to the linguistic example in Ref. [20], this suggests for our disease networks that the incorporation of space into the null model accounts for the majority of the structure present in the network. The spatial structure that we removed likely includes the structure that corresponds to the climate variation that causes different epidemic patterns in the jungle, coastal, and mountainous provinces. The only variable that we were able to identify as influencing community structure when using spatial null models is province population: the highly populated (and typically coastal) provinces forming singleton communities. These highly populated provinces are likely to be economic centers, with many people traveling there from the other provinces and thereby transmitting the disease [57, 74, 93, 94].

These provinces could then be the seeds of epidemics for the other coastal and mountainous provinces, and two studies have in fact reported (so-called) “hierarchical” transmission of dengue from populous regions to those with low populations in both Peru and Thailand [64, 95]. This situation could lead to high correlations across atypically long distances compared with the majority of the data, which could in turn cause populous provinces to be assigned to singleton communities. Additionally, it is known that population size influences dengue transmission: the basic reproductive number *R*_0_ and disease persistence (i.e., the fraction of weeks with disease cases) are positively correlated with population size, and the attack rates are negatively correlated with it [64, 67].

The incorporation of spatial information into null models for community detection is both interesting and desirable. As we have illustrated in the present paper, however, there are many nuances that it is important to consider. We have also demonstrated that it is important to develop null models that incorporate generative mechanisms for human mobility and flux. We similarly expect that domain-dependent, mechanistic null models will also be crucial in many other applications.

## Acknowledgements

MS was supported by the EPSRC Systems Biology Doctoral Training Centre. GC acknowledges support from grant number 1R01GM100471-01 from the National Institute of General Medical Sciences (NIGMS) at the National Institutes of Health and the Multinational Influenza Seasonal Mortality Study (MISMS) led by the Fogarty International Center, National Institutes of Health [96]. MAP was supported by the European Commission FET-Proactive project PLEXMATH (Grant No. 317614) and a grant (EP/J001759/1) from the EPSRC.

We thank Marya Bazzi, Vittoria Colizza, Andrew Elliott, Lucas Jeub, Chiara Poletto, Heather Harrington, and Felix Reed-Tsochas for helpful discussions. We also thank Andrew Elliott for reading and commenting on this manuscript and help with creating the figures. The dengue fever data was collected by the Peruvian Ministry of Health [97]. We obtained province locations from the Geonames.org website [89], and we obtained their populations from the Peruvian Instituto Nacional de Estadística e Informática (INEI) [75].

## A. Spatial benchmarks: Variation of Information

Normalized variation of information (NVI) [62] is a viable alternative similarity measure to NMI for the spatial benchmark networks. In contrast to NMI, variation of information (VI) and NVI are metrics in the mathematical sense. Both measures are related to mutual information. For a partition *X* = {*X*_1_, *X*_2_, … *X_K_* } with *K* communities, VI is defined as

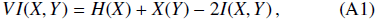

where 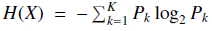 is the entropy of the random variable associated to partition X, the quantity 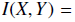 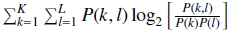 is the mutual information, *P(k)* and *P*(*l*) are the respective marginal probabilities of observing communities *k* and *l* in partitions *X* and *Y*, and *P*(*k, l*) is the joint probability of observing communities *k* and *l* simultaneously in partitions *X* and *Y*). VI is equal to 0 if partitions *X* and *Y* are identical, and *VI* (*X, Y*) < log_2_ *N*, where *N* is the number of nodes in the whole network. Normalizing VI yields NVI, which is given by [63]

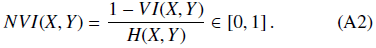

See Refs. [62, 63] for additional discussions. As one can see in Fig. 17, both NMI and NVI perform similarly and neither gives visibly better precision.

**FIG. 17.**
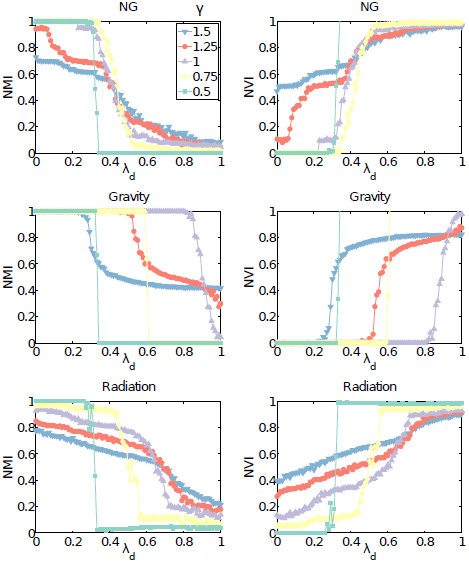
(Left) Normalized mutual information (NMI) and (right) normalized variation of information (NVI) between algorithmically-detected partitions, which we obtain by maximizing modularity, and planted partitions in the uniform population distance static spatial benchmarks with *N* = 50 cities, a grid size of *l* = 10, and a density parameter of *μ* = 50. We examine the partitions for different values of the resolution parameter *γ* as a function of inter-community connectivity *λ_d_* using the (top) NG null model, (middle) gravity null model, and (bottom) radiation null model.

## B. Spatial benchmarks: Variation of the number of cities *N*

We now vary the number *N* of cities in benchmarks with a fixed size of *l* = 10, density parameter of *μ* = 100, and a uniform population of 100 people in each city. In Fig. 18, we plot the NMI of algorithmic partitions versus planted partitions for several values of the resolution parameter *γ* using the NG null model and both spatial null models. In combination with Fig. 3 in the main text, which has *N* = 50 cities, we observe no qualitative changes in NMI aside from an expected increase in variability when *N* is small.

**FIG. 18.**
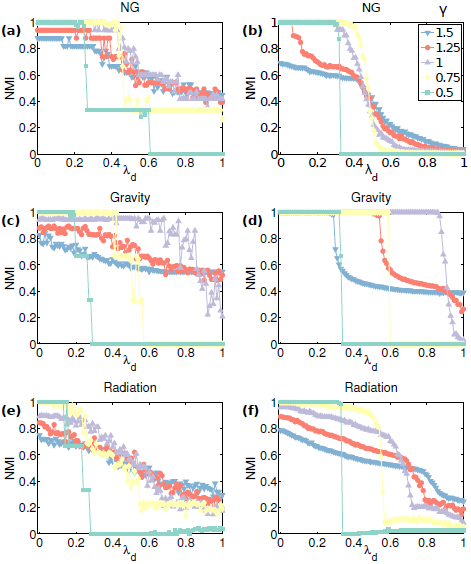
Uniform population static benchmarks: NMI scores between algorithmically detected partitions, which we obtain by maximizing modularity, and planted partitions in static uniform population distance benchmarks with *l* = 10, a density parameter of *μ =* 100, and uniform populations of 100 for different numbers of cities in an underlying space of the same size. The number of cities is (left) *N* = 10, and (right) *N* = 90. We use the NG (top), gravity (middle), and (bottom) radiation null models. See Fig. 3 in the main text for plots with *N* = 50.

## C. Variation of Edge Density Parameter *μ*

We present the results of varying the edge density parameter *μ* in static benchmarks. The edge density has a strong effect on the ability of the modularity-maximization methods to detect communities. For *μ* 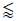 5, we obtain smaller NMI scores than the maximum attained for each particular *λ_d_* for larger *μ* values. (See Figs. 19 and 20.) We therefore focus on using a density parameter of *μ* = 100 in the main text to follow the choice that was used for the benchmarks networks in Ref. [20].

**FIG. 19.**
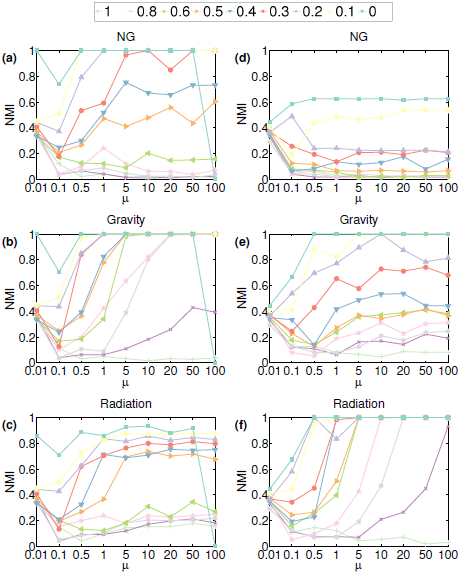
NMI between algorithmically-detected partitions, which we obtain by maximizing modularity with *γ* = 1, and planted partitions for uniform population static spatial benchmarks with *N* = 50, a size parameter of *l* = 10, uniform city populations of 100, and several values of inter-community connectivity *λ_d_*. We plot the NMI scores as a function of the edge density parameter *μ* for (left) the distance benchmark and (right) the flux benchmark.

## D. “Distance and Population” Benchmark

In this section, we construct a “distance and population” spatial benchmark. In Fig. 3 in the main text we observed that the gravity null model performs best on the uniform population distance benchmark, but the NG null model performs better than spatial null models on the random population distance benchmark because the edge placement in that benchmark does not include population information. Here, we study the effects of incorporating population into edge probabilities for the “distance and population” benchmark.

We construct the new type of benchmark network in the same manner as the distance benchmark in Section III, but we now incorporate population into the edge-placement probability by taking 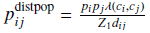. As expected, this brings back the advantage that the gravity null model has for the uniform population distance benchmark (compare Fig. 21 with Fig. 3 in the main text). The radiation null model has the second-best performance on this benchmark, with a better performance than on the random population distance benchmark. However, it does not do as well as it did on the random population flux benchmark (see Fig. 3).

**FIG. 20.**
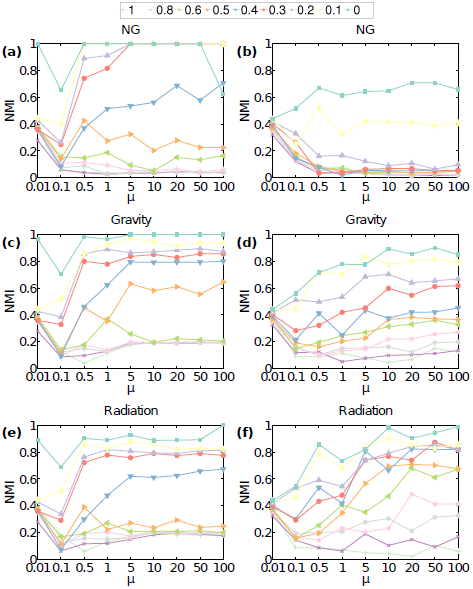
NMI between algorithmically-detected partitions, which we obtain by maximizing modularity with *γ* = 1, and planted partitions for random population static spatial benchmarks with *N* = 50, a size parameter of *l* = 10, city populations *n* selected uniformly at random from [0, 100], and several values of inter-community connectivity *λ_d_*. We plot the NMI scores as a function of the edge density parameter *μ* for (left) the distance benchmark and (right) the flux benchmark.

**FIG. 21.**
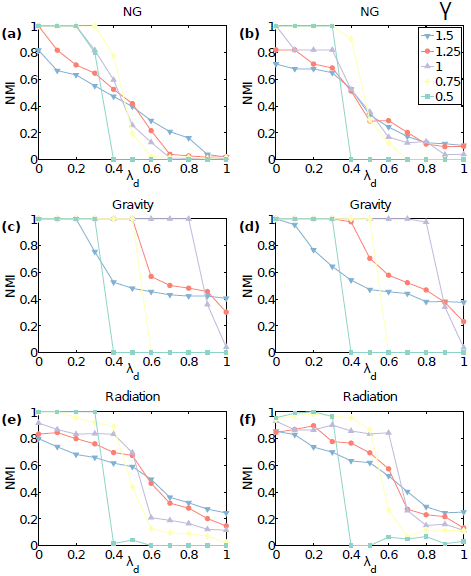
NMI between algorithmically-detected community structure, which we obtain by maximizing modularity, and planted community structure in “distance and population” static spatial benchmarks with (left) uniform populations and (right) random populations. We use *N* = 50, *l* = 10, *m* = 10, *μ* = 100, and *γ* = 1 for various values of *ω* (colored curves) as a function of *λ_d_*. We detected communities by optimizing modularity using the (top) NG, (middle) gravity, and (bottom) radiation null models

## E. Community detection on random population multilayer spatial benchmarks

We now study the influence of the parameters *γ* and *ω* on the community structure for random-population multilayer temporally-stable benchmarks. We first compare the results to our findings from static benchmarks by varying *γ* and *λ_d_* for fixed values of *ω*. We study the performance of modularity maximization with the NG, gravity, and radiation null models on random population benchmarks (see Fig. 4) with parameter values of *N* = 50, *l* = 10, and *m* = 10 layers using *γ* ∈ {0.5, 0.75, 1, 1.25, 1.5} and *ω* ∈ {0.1, 0.25, 0.5, 0.75, 1}. We only show plots for *ω* = 0.1, as the values of w do not noticeably influence the results.

We obtain results that are similar to our results for the corresponding static benchmarks inn Fig. 3.

Once again, we find that the choice of *γ* has a large influence on the quality of algorithmic partitions, and (as with our findings for static benchmarks) that *γ* = 1 seems to yield the best performance (i.e., the largest NMI scores) for low values of *λ_d_*, whereas larger values of *γ* perform better for larger *λ_d_* (see Fig. 22). The effect of varying *γ* is most pronounced for the radiation null model on flux benchmarks.

**FIG. 22.**
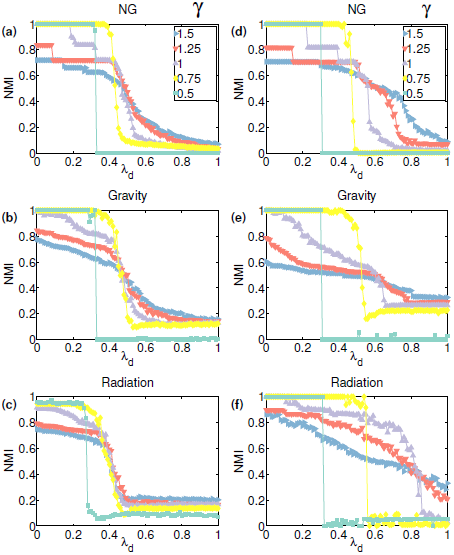
NMI between algorithmically-detected community structure, which we obtain by maximizing modularity, and planted community structure in random-population, temporally-stable multilayer spatial benchmarks. We choose the population of each of the *N* = 50 cities uniformly at random from the set {1,…,100}. We consider various values of the resolution parameter *γ*, and the other parameter values are *l* = 10, *m =* 10, *μ =* 100, and *ω* = 0.1. We plot NMI as a function of *λ_d_* for (left) the distance benchmark and (right) the flux benchmark using the (top) NG, (middle) gravity, and (bottom) radiation null models.

We now examine the NMI of algorithmic versus planted partitions in temporally-stable multilayer benchmarks for fixed *γ* = 1 while varying *ω* and *λ_d_*. As we show in Fig. 23, we find that the value of *ω* usually has very little effect on our ability to detect the planted communities via modularity maximization — the same as for the uniform population temporally stable multilayer benchmarks (see Fig. 5). The parameter *ω* becomes important for the random-population, temporally-evolving multilayer benchmarks in the same manner as what we observed in the main text for uniform population benchmark networks (not shown; see Fig. 6 in the main text for the uniform population results).

**FIG. 23.**
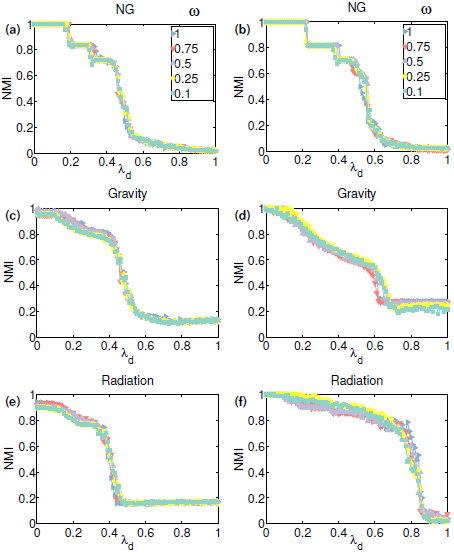
NMI between algorithmically-detected community structure, which we obtain by maximizing modularity, and planted community structure in random-population, temporally-stable multilayer spatial benchmarks. We choose the population of each of the *N* = 50 cities uniformly at random from the set {1,…,100}. We consider various values of the parameter *ω*, and the other parameter values are *l =* 10, *m =* 10, *μ =* 100, and *γ* = 1. We plot NMI as a function of *λ_d_* for (left) the distance benchmark and (right) the flux benchmark using the (top) NG, (middle) gravity, and (bottom) radiation null models.

## F. Province-level Communities for Multilayer Benchmarks

In Fig. 24, we present our results for province-level community detection on uniform population temporally stable multilayer benchmarks. As one can see by comparing these results to those in Fig. 4, we obtain similar NMI scores for the performance of community detection for province-level communities as we did for the ordinary community detection in multilayer networks.

**FIG. 24.**
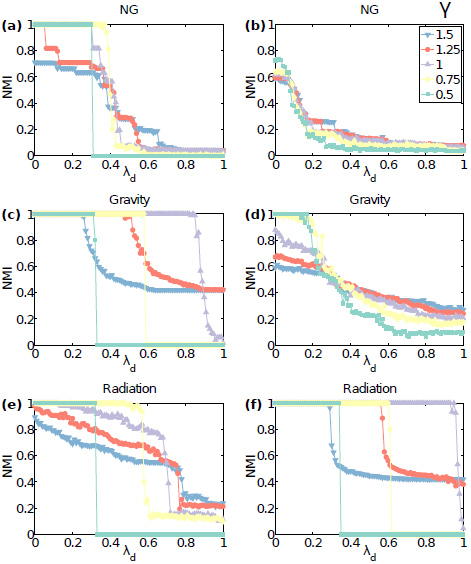
NMI between algorithmically-detected province-level community structures, which we obtain by maximizing modularity, for uniform population (*n_i_* = 100 for all *i*) temporally stable multilayer spatial benchmarks with *m* = 10 layers. Each layer has a single-layer planted partition with *N* = 50 cities, a size parameter of *l* = 10, and a density parameter of *μ =* 100. We use *ω* = 0.1 and consider various values of the resolution parameter *γ*, and we plot NMI as a function of the inter-community connectivity *λ_d_* for (left) the distance benchmark and (right) the flux benchmark.

## G. Community Detection Using a Correlation Null Model with Different values of Δ on dengue fever Data

We now present results of modularity maximization using the correlation null model on correlation networks that we construct from the dengue fever times series with various values of the time-window width Δ ∈ {26, 52, 130, 150} (see Fig. 25). We select the structures that score the highest in comparison to the detailed climate partitions, as in the main text. We observe for small time windows (Δ = 26 and Δ = 52) that the jungle provinces split into two communities. For large time windows (Δ = 130 and Δ = 150}), we obtain very similar results to those that we showed for Δ = 104 in Fig. 13 in the main text. The community structure is dominated by one large community of jungle provinces.

**FIG. 25.**
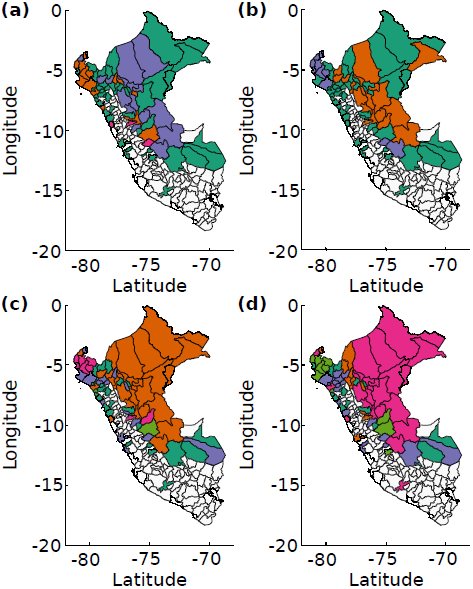
Algorithmically-detected community structures, which we obtain using modularity maximization, for a correlation null model and *γ* = 1 on the sets of static dengue fever correlation networks that we construct using different time-window widths Δ. We show results for (a) Δ = 26, *γ* = 0.1, and layer 532; (b) Δ = 52, *γ* = 0.6, and layer 72; (c) Δ = 130, *γ* = 2.5, and layer 460; and (d) Δ = 150, *γ* = 1, and layer 492. These structures were the highest scoring against the detailed climate partition. We show partitions on a map of Peru and color provinces according to their community. White provinces are ones in which our data does not include any reported cases of dengue fever in the indicated time window.

## H. Community Detection on Aggregated dengue fever Data

In Fig. 26, we show additional results of community detection on fully aggregated networks (i.e., we use *τ* = 1 and Δ = 779) from the dengue fever times series. In Section IV D 1 of the main text, in Fig. 15(a) we showed the results of modularity maximization using the NG null model. We now also show similar results for the gravity, radiation, and correlation null models. The gravity, radiation, and correlation null models find one large community and a few small communities. Because of the aggregation, we have lost the rich set of information that we were able to study using multilayer community detection.

**FIG. 26.**
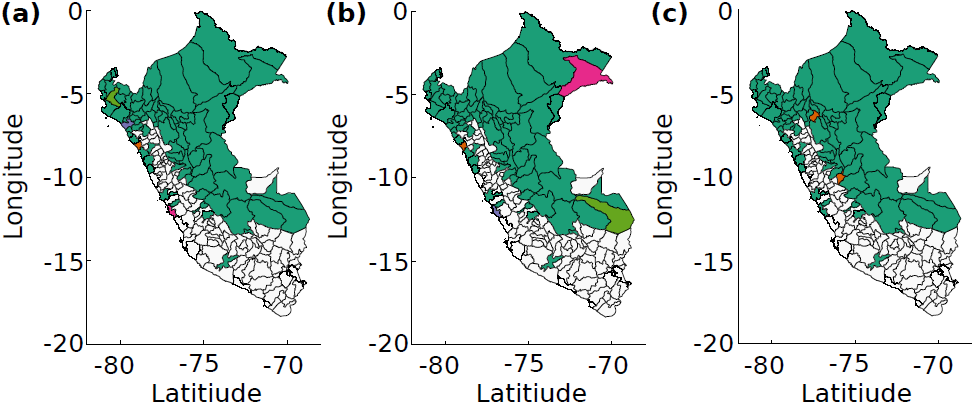
Algorithmically-detected community structures, which we obtain using modularity maximization, for static dengue fever correlation networks that we construct using the entire set of time series (i.e., we use *τ* = 1 and Δ = 779) using (a) the gravity null model, (b) the radiation null model, and (c) the correlation null model for a resolution-parameter value of *γ* = 1. We color provinces on a map of Peru according to their community assignments. White provinces are ones in which our data does not include any reported cases of dengue fever in the indicated time window.

## Footnotes

1 Although the directionality of fluxes is an important factor to study, we wish to keep our null models as simple as possible in order to focus on the effect of incorporating space into them. Additionally (and again for simplicity), we will construct our disease-correlation networks using Pearson correlations, so we will study the community structure of undirected networks. If one instead constructs a directed networks—e.g., by including a time delay when measuring the similarity of time series, considering ideas such as Granger causality, or otherwise measuring similarity in a way that produces a directed network (see, e.g., Ref. [59]), then it would also be desirable to construct a directed version of the radiation null model. Clearly, this is an interesting future direction, but it is beyond the scope of our study.

2 We discuss the *correlation null model*, which was recently introduced in [18], in Section IV D.

3 Reference [59] compared several methods to calculate similarity networks from time-series data. Our focus in the present paper is on generalizing and evaluating null models, so we use Pearson correlations for simplicity.

4 See an analogous discussion of time-window choice in Ref. [82] in the context of financial networks.

## References

[1] M. E. J. Newman, Networks: An Introduction (Oxford University Press, 2010).

[2] S. Wasserman and K. Faust, Social Network Analysis: Methods and Applications (Cambridge University Press, 1994).

[3] P. Holme and J. Saramäki, Phys. Rep. 519, 97 (2012).

[4] P. Holme and J. Saramäki, eds., Temporal Networks (Springer-Verlag, 2013).

[5] M. Kivelä, A. Arenas, M. Barthelemy, J. P. Gleeson, Y. Moreno, and M. A. Porter, “Multilayer Networks,” (2014), arXiv:1309.7233.

[6] S. Boccaletti, G. Bianconi, R. Criado, C. I. del Genio, J. Gómez-Gardeñes, M. Romance, I. Sendiña-Nadal, Z. Wang, and M. Zanin, “The structure and dynamics of multilayer networks,” (2014), arXiv:1407.0742 (Physics Reports, in press, doi:10.1016/j.physrep.2014.07.001).

[7] M. Barthélemy, Phys. Rep. 499, 1 (2011).

[8] S. Fortunato, Phys. Rep. 486, 75 (2010).

[9] M. A. Porter, J.-P. Onnela, and P. J. Mucha, Notices Amer. Math. Soc. 56, 1082 (2009).

[10] A. L. Traud, P. J. Mucha, and M. A. Porter, Physica A 391, 4165 (2012).

[11] A. S. Waugh, L. Pei, J. H. Fowler, P. J. Mucha, and M. A. Porter, “Party polarization in Congress: A network science approach,” (2012), arXiv:0907.3509.

[12] A. C. F. Lewis, N. S. Jones, M. A. Porter, and C. M. Deane, BMCSysBio l4 (2010).

[13] V. Spirin and L. A. Mirny, Proc. Natl. Acad. Sci. USA 100, 12123 (2003).

[14] M. Newman and M. Girvan, Phys. Rev. E69, 026113 (2004).

[15] M. E. J. Newman, Phys. Rev. E74, 036104 (2006).

[16] R. Lambiotte, J.-C. Delvenne, and M. Barahona, “Laplacian dynamics and multiscale modular structure in networks,” (2009), arXiv:0812.1770.

[17] B. H. Good, Y. A. de Montjoye, and A. Clauset, Phys. Rev. E 81, 046106 (2010).

[18] M. MacMahon and D. Garlaschelli, “Unbiased community detection for correlation matrices,” (2013), arXiv:1311.1924.

[19] M. Bazzi, M. A. Porter, M. McDonald, D. J. Fenn, S. Williams, and S. D. Howison, “Community structure in multilayer networks of financial-asset correlations,” (2014), in preparation.

[20] P. Expert, T. S. Evans, V. D. Blondel, and R. Lambiotte, Proc. Natl. Acad. Sci. USA 108, 7663 (2011).

[21] T. Y. Berger-Wolf and J. Saia, in KDD ‘06: Proceedings of the 12th ACM S/GKDD international conference on Knowledge discovery and data mining (2006) pp. 523–528.

[22] D. J. Fenn, M. A. Porter, M. McDonald, S. Williams, N. F. Johnson, and N.S. Jones Chaos, 19, 033119 (2009).

[23] S.-Y. Chan, P. Hui, and K. Xu, in Complex Sciences, Lecture Notes of the Institute for Computer Sciences, Social Informatics and Telecommunications Engineering, Vol. 4, edited by J. Zhou (Springer Berlin Heidelberg, 2009) pp. 1154–1159.

[24] P. J. Mucha, T. Richardson, K. Macon, M. A. Porter, and J.-P. Onnela, Science 328, 876 (2010).

[25] V. Kawadia and S. Sreenivasan, Scientific Reports 2, 794 (2012).

[26] Y. Chen, V. Kawadia, and R. Urgaonkar, “Detecting Overlapping Temporal Community Structure in Time-Evolving Networks,” (2013), arXiv:1303.7226.

[27] D. S. Bassett, M. A. Porter, N. F. Wymbs, S. T. Grafton, J. M. Carlson, and P. J. Mucha, Chaos 23, 013142 (2013).

[28] F. Cerina, V. De Leo, M. Barthelemy, and A. Chessa, PLoS One 7, e37507 (2012).

[29] J. Hannigan, G. Hernandez, R. M. Medina, P. Roos, and P. Shakarian, “Mining for Spatially-Near Communities in Geo-Located Social Networks,” (2013), arXiv:1311.1924.

[30] P. Shakarian, P. Roos, D. Callahan, and C. Kirk, “Mining for Geographically Disperse Communities in Social Networks by Leveraging Distance Modularity,” (2013), arXiv:1305.3668.

[31] D. S. Bassett, E. T. Owens, M. A. Porter, M. L. Manning, and K. E. Daniels, “Practical methods for the examination of force chain network architecture in granular materials,” (2014), in preparation.

[32] A. Barrat, M. Barthelemy, and A. Vespignani, Dynamical Processes on Complex Networks (Cambridge University Press, 2008).

[33] D. Balcan, H. Hu, B. Goncalves, P. Bajardi, C. Poletto, J. J. Ramasco, D. Paolotti, N. Perra, M. Tizzoni, W. Van den Broeck, V. Colizza, and A. Vespignani, BMC Medicine 7, 45 (2009).

[34] V. Colizza, R. Pastor-Satorras, and A. Vespignani, Nature Physics 3, 276 (2007).

[35] Y. Xia, O. N. Bjornstad, and B. T. Grenfell, Am. Nat. 164, 267 (2004).

[36] M. De Domenico, A. Solé-Ribalta, E. Cozzo, M. Kivelä, Y. Moreno, M. A. Porter, S. Gómez, and A. Arenas, Phys. Rev. X3, 041022 (2013).

[37] M. E. J. Newman, Phys. Rev. E 70, 056131 (2004).

[38] J. Reichardt and S. Bornholdt, Phys. Rev. E74, 016110 (2006).

[39] J.-P. Onnela, D. J. Fenn, S. Reid, M. A. Porter, P. J. Mucha, M. D. Fricker, and N. S. Jones, Phys. Rev. E 86 (2012).

[40] U. Brandes, D. Delling, M. Gaertler, R. Gorke, M. Hoefer, Z. Nikoloski, and D. Wagner, IEEE Transactions on Knowledge and Data Engineering 20, 172 (2008).

[41] V. D. Blondel, J.-L. Guillaume, R. Lambiotte, and E. Lefebvre, J. Stat. Mech. 2008, P10008 (2008), 0803.0476v2.

[42] I. S. Jutla, P. J. Mucha, and L. Jeub, “GenLouvain: A generalized Louvain method for community detection implemented in Matlab,” (2011–2012), http://netwiki.amath.unc.edu/ GenLouvain.

[43] A. Lancichinetti and S. Fortunato, Sci Rep 2, 794 (2012).

[44] M. E. J. Newman, Proc. Natl. Acad. Sci. US A103, 8577 (2006).

[45] M. E. J. Newman, Phys. Rev. E 74 (2006).

[46] G. K. Zipf, Am. Sociol. Rev. 11, 677 (1946).

[47] J. Q. Stewart, Geogr. Rev. 37, 461 (1947).

[48] J. Q. Stewart and W. Warntz, J. Reg. Sci. 1, 99 (1958).

[49] A. G. Wilson, Transport. Res. 1, 253 (1958).

[50] D. Balcan, V. Colizza, B. Goncalves, H. Hu, J. J. Ramasco, and Vespignani, Proc. Natl. Acad. Sci. USA 106, 21484 (2009).

[51] S. H. Lee, R. Ffrancon, D. M. Abrams, B. J. Kim, and M. A. Porter, “ Matchmaker, Matchmaker, Make Me a Match: Migration of Populations Via Marriages in the Past,” (2014), arXiv:1310.7532.

[52] A.-C. Disdier and K. Head, Rev Econ. Stat. 90, 37 (2008).

[53] K. E. Jones, N. G. Patel, M. A. Levy, A. Storeygard, D. Balk, J. Gittleman, and P. Daszak, Nature 451, 990 (2008).

[54] F. Gonzalez, M. C. Maritan, A. Simini, and A.-L. Barabasi, Nature 484, 96 (2012).

[55] S. Goh, K. Lee, J. S. Park, and M. Y. Choi, Phys. Rev. E 86, 026102 (2012).

[56] A. P. Masucci, J. Serras, A. Johansson, and M. Batty, Phys. Rev. E88, 022812 (2013).

[57] S. T. Stoddard, A. Morrison, G. M. Vazquez-Prokopec, V. Paz Soldan, T. J. Kochel, U. Kitron, J. Elder, and T. W. Scott, PLoS Negl. Trop. Dis. 3, e481 (2009).

[58] F. Simini, A. Maritan, and Z. Néda, PLoS One 8, e60069 (2013).

[59] S. M. Smith, K. L. Miller, G. Salimi-Khorshidi, M. Webster, C. F. Beckmann, T. E. Nichols, J. D. Ramsey, and M. W. Woolrich, NeuroImage 54, 875 (2011).

[60] A. Strehl, J. Ghosh, and C. Cardie, J. Mach. Learn. Res. 3, 583 (2002).

[61] A. L. Traud, E. D. Kelsic, P. J. Mucha, and M. A. Porter, SIAM Review 53, 526 (2011).

[62] M. Meilá, J. Multivariate Anal. 98, 873 (2007).

[63] A. Kraskov, H. Stägbauer, R. G. Andrzejak, and P. Grassberger, Europhys. Lett. 70, 278 (2005).

[64] G. Chowell, C. A. Torre, C. Munáyco-Escate, L. Sudrez-Ognio, R. López-Cruz, J. M. Hyman, and C. Castillo-Chavez, Epidemiol. Infect. 136, 1667 (2008).

[65] M. G. Guzman, S. B. Halstead, H. Artsob, P. Buchy, J. Farrar, D. J. Gubler, E. Hunsperger, A. Kroeger, H. S. Margolis, E. Martínez, M. B. Nathan, J. Pelegrino, C. Simmons, S. Yoksan, and R. W. Peeling, Nat. Rev. Microbiol. 8, S7 (2010).

[66] M. Guzman, J. Clin. Virol. 27, 1 (2003).

[67] G. Chowell, B. Cazelles, H. Broutin, and C. V. Munayco, BMC InfectiousDiseases11, 164 (2011).

[68] M. A. Johansson, F. Dominici, and G. E. Glass, PLoS Negl. Trop. Dis. 3, e382 (2009).

[69] T. H. Jetten and D. A. Focks, Am. J. Trop. Med. Hyg. 57, 285 (1997).

[70] C. F. Li, T. W. Lim, L. L. Han, and R. Fang, Southeast Asian J. Trop. Med. Public Health 16, 560 (1985).

[71] C. Depradine and E. Lovell, Int. J. Environ. Health Res. 6, 429 (2004).

[72] J. Keating, Soc. Sci. Med. 53, 1587 (2001).

[73] G. Chowell and F. Sanchez, J. Environ. Health 68, 40 (2006).

[74] L. C. Harrington, T. W. Scott, K. Lerdthusnee, R. C. Coleman, A. Costero, G. G. Clark, J. J. Jones, S. Kitthawee, P. Kittayapong, R. Sithiprasasna, et al., Am. J. Trop. Med. Hyg. 72, 209 (2005).

[75] Instituto Nacional de Estadística e Informática (INEI), http://www.inei.gob.pe/.

[76] T. J. Kochel, D. M. Watts, S. B. Halstead, C. G. Hayes, A. Espinoza, V. Felices, R. Caceda, C. T. Bautista, Y. Montoya, S. Douglas, and K. L. Russell, Lancet 360, 310 (2002).

[77] Y. Montoya, S. Holechek, O. Caceres, A. Palacios, J. Burans, C. Guevara, F. Quintana, V. Herrera, E. Pozo, and E. Anaya, Dengue Bulletin 27, 52 (2003).

[78] L. S. Lloyd, “Best practices for dengue prevention and control in the Americas. Strategic report, Environmental Health Project Office of Health Infectious Diseases and Nutrition,” (2003).

[79] D. S. Bassett, N. F. Wymbs, M. A. Porter, P. J. Mucha, J. M. Carlson, and S. T. Grafton, Proc. Natl. Acad. Sci. USA 108, 7641 (2011).

[80] P. J. Mucha and M. A. Porter, Chaos 20, 041108 (2010).

[81] K. T. Macon, P. J. Mucha, and M. A. Porter, Physica A 391, 343 (2012).

[82] D. J. Fenn, M. A. Porter, P. J. Mucha, M. Mcdonald, S. Williams, N. F. Johnson, and N. S. Jones, Quantitative Finance, 1 (2012).

[83] A. J. Tatem, D. J. Rogers, and S. I. Hay, in Global Mapping of Infectious Diseases: Methods, Examples and Emerging Applications, Advances in Parasitology, Vol. 62, edited by A. G. S I. Hay and D. J. Rogers (Academic Press, 2006) pp. 293–343.

[84] J. Truscott and N. M. Ferguson, PLoS Comput Biol 8, e1002699 (2012).

[85] L. Anselin, Geogr. Anal. 27, 93 (1995).

[86] H. J. Wearing and P. Rohani, Proc. Natl. Acad. Sci. USA 103, 11802 (2006).

[87] C. Thiemann, F. Theis, D. Grady, R. Brune, and D. Brockmann, PLoS One 5, e15422 (2010).

[88] C. Ratti, S. Sobolevsky, F. Calabrese, C. Andris, J. Reades, M. Martino, R. Claxton, and S. H. Strogatz, PLoS One 5, e14248 (2010).

[89] Geonames.org, http://www.geonames.org/.

[90] G. Kuno, in Dengue and Dengue Hemorrhagic Fever., edited by K. G. Gubler D J (CAB International, Wallingford, UK, 1997) pp. 61–88.

[91] M. Mehta, Random Matrices, 3rd ed. (Academic Press, San Diego, California, 2004).

[92] S. Sinha, A. Chatterjee, A. Chakraborti, and B. K. Chakrabarti, Econophysics (Wiley-VCH, 2011).

[93] L. Osorio, J. Todd, and D. J. Bradley, Am. J. Trop. Med. Hyg. 71, 380 (2004).

[94] P. Martens and L. Hall, Emerging Infect Dis., 6 (2000).

[95] D. A. T. Cummings, R. A. Irizarry, N. E. Huang, T. P. Endy, A. Nisalak, K. Ungchusak, and D. S. Burke, Nature 427, 344 (2004).

[96] Fogarty International Center, National Institutes of Health, http://www.origem.info/misms/index.php.

[97] Directiòn General de Epidemiologia, http://www.dge.gob.pe.

